# Kazrin is an endosomal adaptor for dynein/dynactin

**DOI:** 10.1101/2021.08.30.458243

**Authors:** Ines Hernandez-Perez, Adrian Baumann, Javier Rubio, Henrique Girao, Elena Rebollo, Anna M. Aragay, Maria Isabel Geli

## Abstract

Kazrin is a protein widely expressed in vertebrates whose depletion causes a myriad of developmental defects, in part derived from altered cell adhesion, impaired cell migration and failure to undergo Epidermal to Mesenchymal Transition (EMT). However, the primary molecular role of kazrin, which might contribute to all these functions, has not been elucidated yet. We previously identified one of its isoforms, kazrin C, as a protein that potently inhibits clathrin-mediated endocytosis when overexpressed. We now generated kazrin knock out Mouse Embryonic Fibroblasts (MEFs) to investigate its endocytic function. We found that kazrin depletion delays perinuclear enrichment of internalized material, indicating a role in endocytic traffic from Early (EE) to Recycling Endosomes (REs). Consistently, we found that the C-terminal domain of kazrin C, predicted to be an Intrinsically Disordered Region (IDR), directly interacts with several components of the EEs, and that kazrin depletion impairs centripetal motility of EEs. Further, we noticed that the N-terminus of kazrin C shares homology with dynein/dynactin adaptors and that it directly interacts with the dynactin complex and the dynein Light Intermediate Chain 1 (LIC1). Altogether, the data indicate that one of the primary kazrin functions is to facilitate endocytic recycling via the perinuclear endocytic compartment, by promoting microtubule and dynein/dynactin-dependent transport of EEs or EE-derived transport intermediates to the RE.

## INTRODUCTION

Kazrin is a highly conserved and broadly expressed vertebrate protein, which was first identified as a transcript present in human brain (Kikuno et al., 1999). The human kazrin gene is located on chromosome 1 (1p36.21) and encodes at least seven isoforms (A-F and K), generated by alternative splicing (Groot et al., 2004; Nachat et al., 2009; Wang et al., 2009). From those, kazrin C is the shorter isoform that constitutes the core of all other versions, which bear N or C-terminal extensions. Since its discovery, several laboratories have reported a broad range of roles for the different kazrin isoforms in a myriad of experimental model systems. Thus, in humans, kazrin participates in structuring the skin cornified envelope and it promotes keratinocyte terminal differentiation (Groot et al., 2004; Sevilla et al., 2008a). In U373MG human astrocytoma cells, kazrin depletion leads to caspase activation and apoptosis (Wang et al., 2009). In *Xenopus* embryos, kazrin is important to maintain the ectoderm integrity and its depletion also causes skin blisters, probably derived from defects in establishing cell-cell contacts (Sevilla et al., 2008b). In addition, depletion of kazrin causes craniofacial development defects, linked to altered EMT and impaired migration of neural crest cells (Cho et al., 2011). The subcellular localization of kazrin recapitulates its functional diversity. Depending on the isoform and cell type under analysis, kazrin has been reported to associate with desmosomes (Groot et al., 2004), *adherens* junction components (Cho et al., 2010; Sevilla et al., 2008a), the nucleus (Groot et al., 2004; Sevilla et al., 2008a), or the microtubule cytoskeleton (Nachat et al., 2009). At the molecular level, the N-terminus of kazrin, predicted to form a coiled-coil, directly interacts with several p120-catenin family members (Sevilla et al., 2008b), as well as with the desmosomal component periplakin (Groot et al., 2004), and it somehow regulates RhoA activity (Groot et al., 2004; Sevilla et al., 2008a) (Cho et al., 2010). How kazrin orchestrates such many cellular functions at the molecular level is far from being understood.

We previously isolated human kazrin C as a cDNA that when overexpressed potently inhibits clathrin-mediated endocytosis (Schmelzl and Geli, 2002). In the present work, we generated kazrin knock out (kazKO) MEFs to analyze its role in endocytic traffic in detail. We found that depletion of kazrin caused accumulation of peripheral EEs and delayed transfer of endocytosed transferrin (Tfn) to the perinuclear region, where the REs concentrate. Consequently, cellular functions requiring intact endosomal traffic through the REs, such as cell migration and cytokinetic abscission, were also altered. Consistent with its role in endocytic traffic, we found that the kazrin C C-terminal portion, predicted to be an IDR, interacted with different components of the EEs, it was required to form condensates on those organelles and it was necessary to sustain efficient transport of internalized Tfn to the perinuclear region. Further, the N-terminus of kazrin C, which shared considerable homology with dynein/dynactin adaptors, directly interacted with the dynactin complex and LIC1, and depletion of kazrin impaired the centripetal motility of Tfn-loaded EEs. The data thus suggested that kazrin facilitates transfer of endocytosed material to the pericentriolar RE by acting as a dynein/dynactin adaptor for EEs or EE-derived transport intermediates.

## RESULTS

### Kazrin depletion impairs endosomal traffic

We originally identified kazrin C as a human brain cDNA, whose overexpression causes the accumulation of the Tfn Receptor (TfnR) at the Plasma Membrane (PM) in Cos7 cells (Schmelzl and Geli, 2002), suggesting that kazrin might be involved in clathrin-mediated endocytic uptake from the PM. However, treatment of Cos7 cells with a shRNA directed against kazrin (shkaz) (Fig. S1A) did not inhibit endocytic internalization but it rather increased the accumulation of Texas Red-Tfn (TxR-Tfn) upon a 2 hour exposure to the ligand (Fig. S1B & C), indicating that depletion of kazrin either exacerbated endocytic uptake or inhibited endocytic recycling. The distribution of TxR-Tfn labeled endosomes was also altered in the shkaz treated cells, as compared with that of untreated cells or cells transfected with a control shRNA (shCTR). In WT and shCTR treated cells, TxR-Tfn accumulated in the perinuclear region, where the RE is located, as previously described (Mellman, 1996). In contrast, TxR-Tfn labelled endosomes appeared more scattered todwards the cell periphery in shkaz treated cells (Fig. S1B & C). The accumulation of endocytosed material at the periphery suggested that kazrin might play a post-internalization role in the endocytic pathway, possibly in the transport of material towards the perinuclear RE.

shRNA transfection in Cos7 cells did not achieved complete kazrin depletion in a reproducible manner and it hampered complementation. To overcome these problems, we generated kazrin knockout MEFs (kazKO MEFs) using the CRISPR CAS9 technology and we used a lentiviral system to subsequently create two cell lines that expressed GFP or GFP-kazrin C upon doxycycline induction (Fig. S1D). Immunoblot analysis demonstrated that the expression level of GFP-kazrin C in the absence of doxycycline or upon a short o/n (up to 12 hour) incubation was similar to that of the endogenous kazrin (low expression, 1 to 4 times the endogenous kazrin expression level) (Fig. S1E). Under these conditions, the GFP-kazrin C was barely detectable by fluorescence microscopy (Fig. S1F). This might explain why none of the commercially available or home-made anti-kazrin antibodies detected a specific signal in Wild Type (WT) MEFs. Doxycycline incubation for longer periods (up to 24 hours induction) resulted in moderate expression (4 to 8 times the endogenous kazrin expression levels) (Fig. S1E), but allowed us to clearly visualize its localization by microscopy (Fig. S1F).

To better discern on the possible effects of kazrin depletion on endocytic uptake or in subsequent trafficking events, WT and kazKO cells were exposed to a short, 10 minutes incubation with TxR-Tfn, fixed, and analyzed. Under these experimental conditions, no differences in the amount of internalized TxR-Tfn were observed between WT and kazKO cells (Fig. S1G & H), suggesting that kazrin did not play a relevant role in the formation of endocytic vesicles from the PM, but it might rather work downstream in the pathway. In agreement with this view and with the Cos7 shRNA data, kazKO MEFs accumulated TxR-Tfn in the periphery in a perinuclear enrichment assay, whereas the endocytosed cargo accumulated in the perinuclear region in most WT cells (Fig. 1A & B). Perinuclear accumulation of TxR-Tfn was restored by low, physiological expression of GFP-kazrin C but not GFP (Fig. 1A & B), indicating a direct role of kazrin in the process. No significant difference between the kazKO and the kazKO GFP-expressing cells could be detected in these experiments. Therefore, in order to simplify the experimental design, further assays were normalized to the most accurate isogenic kazKO background, namely the kazKO cells when comparing to the WT, and the kazKO GFP expressing cells when compared to kazKO MEF expressing GFP-kazrin C.

**Figure 1.**
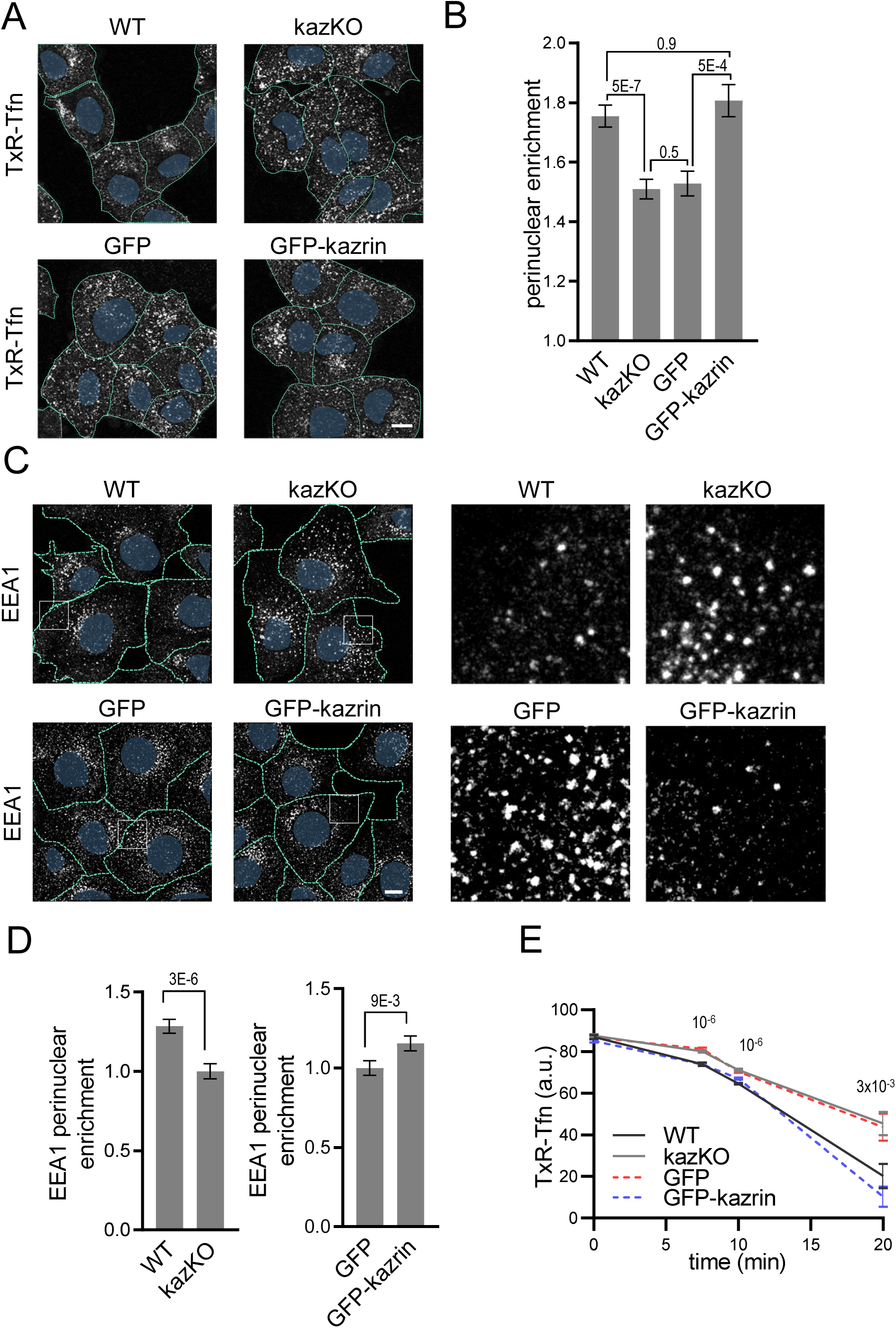
Kazrin depletion impairs endosomal traffic. (**A**) Confocal images of WT and kazKO MEF or kazKO MEF expressing low levels (See M&M) of GFP or GFP-kazrin C incubated with 20 µg/ml TxR-Tfn at 16°C and chased at 37°C for 10 minutes. Scale bar, 10 μm. Cell borders are indicated by dashed lines and the nuclei in blue. (**B**) Mean ± SEM TxR-Tfn perinuclear enrichment for the cells described in A, after 10 minutes incubation with the probe. The mean fluorescence intensity within a circle of 10 µm in the perinuclear region was divided by the mean signal in the cell. p values of two-tailed Mann-Whitney tests are shown. n > 58 for each sample. (**C**) Confocal images of the WT and kazKO MEF, or kazKO MEF expressing GFP or GFP-kazrin C at low levels, fixed and stained with α-EEA1 and A568-conjugated secondary antibodies. Magnified insets showing endosomes in the peripheral areas for each cell type are shown on the right. Scale bar, 10 μm. Cell borders are indicated with dashed lines and the nuclei in blue. (**D**) Mean ± SEM EEA1 perinuclear enrichment. The mean EEA1 fluorescence intensity within a circle of 10 µm in the perinuclear region was quantified and divided by the mean fluorescence intensity in the cell. The values were normalised to the corresponding kazKO cells (either kazKO or kazKO GFP). p values of two-tailed Mann-Whitney tests are shown. n > 80 for each sample. (**E**) Mean ± SEM of TxR-Tfn fluorescence intensity per cell in WT and kazKO MEFs, or kazKO MEFs expressing low levels of GFP and GFP-kazrin C, at the indicated time points after the cells were incubated 30 minutes with 20 µg/ml TxR-Tfn at 16°C to allow accumulation of cargo in EEs, washed and released (time 0) with non-labelled Tfn at 37°C, to allow recycling. Data was normalized to the average intensity at time 0. p values of two-tailed Student t tests are shown. n > 16 per sample and time point.

To evaluate if the scattering of TxR-Tfn endosomes was due a defect in the transfer of material form the EEs to the RE or if it was caused by the dispersal of the RE, we analyzed the distribution of the EE and the RE markers EEA1 (Early Endosome Autoantigen 1) and RAB11 (Ras-Related in Brain 11), respectively. We observed that kazKO MEFs accumulated peripheral, often enlarged, EEA1 positive structures (Fig. 1C & D). The perinuclear distribution of the RE, was however not significantly affected in the knock out cells (Fig. S1I & J). Again, low expression of GFP-kazrin C but not GFP recovered the EEA1 perinuclear distribution (Fig. 1C & D). The data thus suggested that kazrin promotes transfer of endocytosed material towards the perinuclear region, where the RE is located. Consistent with a role of kazrin in endocytic traffic towards the perinuclear RE, recycling of TxR-Tfn back to the PM was diminished in kazKO cells (Fig. 1E), albeit traffic back to the surface was not blocked. A complete block in recycling was not to be expected because, in addition to the RAB11 route, the TfnR can take a RAB4-dependent shortcut to the PM (Sheff et al., 2002). As for the perinuclear Tfn enrichment assays, expression of GFP-kazrin C but not GFP restored the recycling defects installed in the kazKO MEF (Fig. 1E).

To further confirm the specific role of kazrin in endocytic recycling via the perinuclear RE, we analyzed its implication in cellular processes that strongly rely on this pathway, such as cell migration and invasion (Wilson et al., 2018). Analysis of the migration of single WT and kazKO cells through Matrigel demonstrated that depletion of kazrin significantly diminished the migration speed, which, similar to endocytic traffic, was recovered upon re-expressing GFP-kazrin C at low levels, but not GFP (Fig. 2A & B, movie S1). We also observed an increased persistency in kazKO cells (Fig. S2A), but it was not recovered with GFP-kazrin C re-expression (Fig. S2A). Increased persistency might be a secondary effect caused by the trafficking block to the RE, which might accelerate recycling via the shortcut circuit, as previously observed (Perrin et al., 2013; White et al., 2007). The long recycling circuit also plays an important role in the last abscission step during cytokinesis (Pollard and O’Shaughnessy, 2019; Wilson et al., 2005). Consistent with kazrin playing a role in that pathway, kazKO cells had a significant delay in cell separation after cytokinesis, which was again restored by GFP-kazrin C expression (Fig. 2C & D and movie S2).

**Figure 2.**
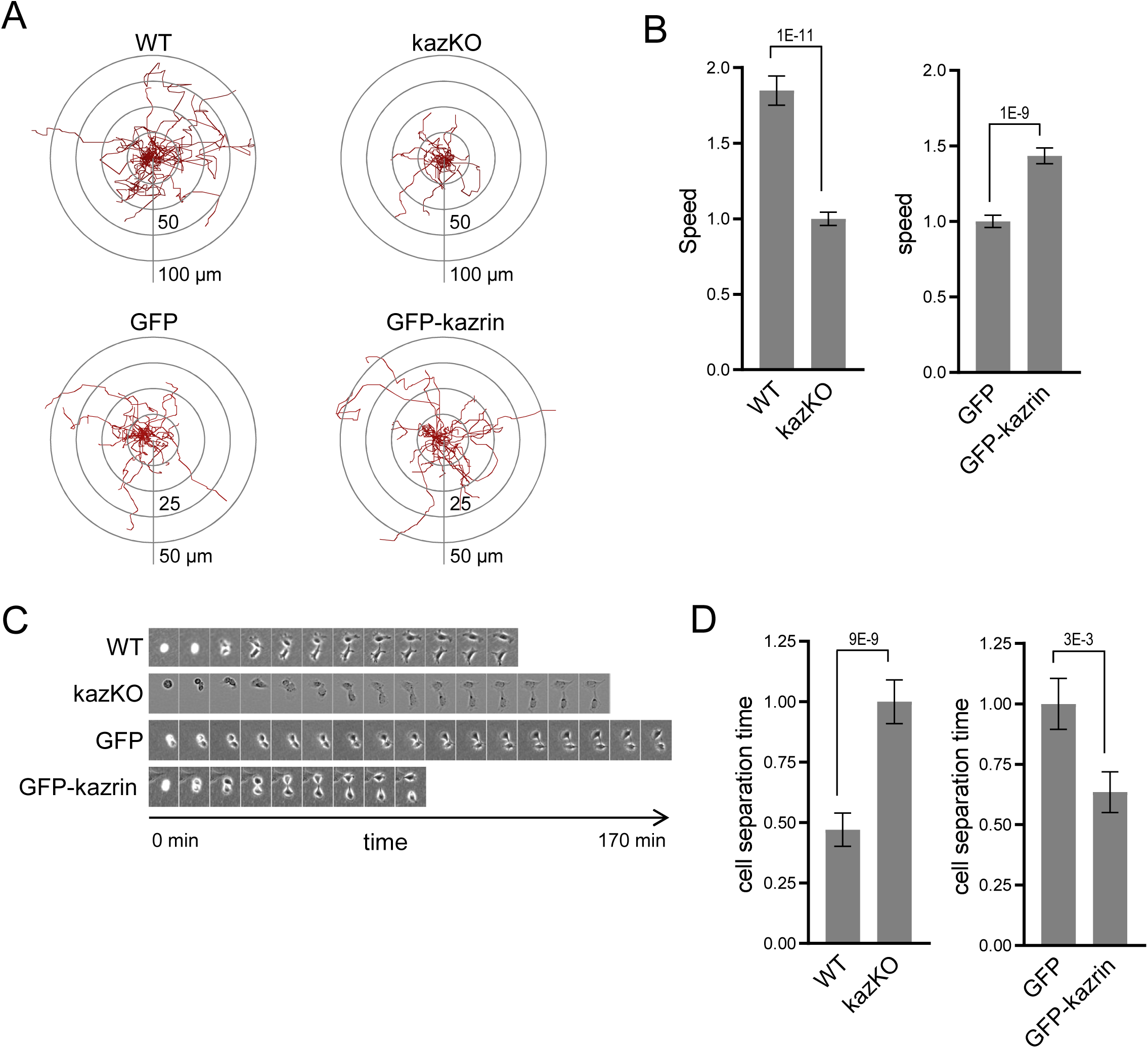
Kazrin depletion impairs cell migration and division. (**A**) Paths described by individually migrating WT and kazKO MEF or kazKO MEFs expressing GFP or GFP-kazrin C at low levels. The cells were embedded in Matrigel and tracked for 9 hours with a 10 minutes time lapse. All tracks start at the (0,0) coordinate in the graph. (**B**) Mean ± SEM speeds of the cells described in (A). The data was normalized to the mean of the corresponding KO cells (either kazKO or kazKO expressing GFP). p values of two-tailed Mann-Whitney tests are shown. n > 100 per condition. (**C**) Time lapse epifluorescence images of WT and kazKO MEFs or kazKO MEFs expressing GFP or GFP-kazrin C at low levels, as they divide. Cells were recorded every 10 minutes. (**D**) Mean ± SEM time lapse between substrate attachment and complete cell separation of the cells described in (C). The data was normalized to the mean of the corresponding KO (kazKO or kazKO expressing GFP). p values of two-tailed Mann-Whitney tests are shown. n > 68 per condition.

### Kazrin is recruited to EEs and directly interacts with components of the endosomal machinery through its C-terminal predicted IDR

Next, we investigated whether endogenous kazrin was present on EEs. For that purpose, we initially used subcellular fractionation and immunoblot because the endogenous protein was not detectable by fluorescence microscopy, nor was GFP-kazrin C expressed at physiological levels. As shown in figure 3A, endogenous kazrin neatly co-fractionated in the lightest fractions with EE markers such as the tethering factor EEA1 (Early Endosome Antigen 1) and EHD (Eps15 Homology Domain) proteins, most likely corresponding to EHD1 and EHD3. On the contrary, it only partially co-fractionated with an early-to-late endosome marker (Vacuolar Protein Sorting 35 ortholog, VPS35) and did not with markers of recycling endosomes (RAB11) or the Golgi apparatus (Golgi Matrix protein 130, GM130) (Fig. 3A). Moderately overexpressed GFP-kazrin C also co-fractionated with EEs, although it appeared slightly more spread towards the RE and Golgi fractions in the gradient (Fig. 3A). Endogenous kazrin localization at EE was confirmed by subcellular fractionation experiments in Cos7 cells (Fig. S2B).

**Figure 3.**
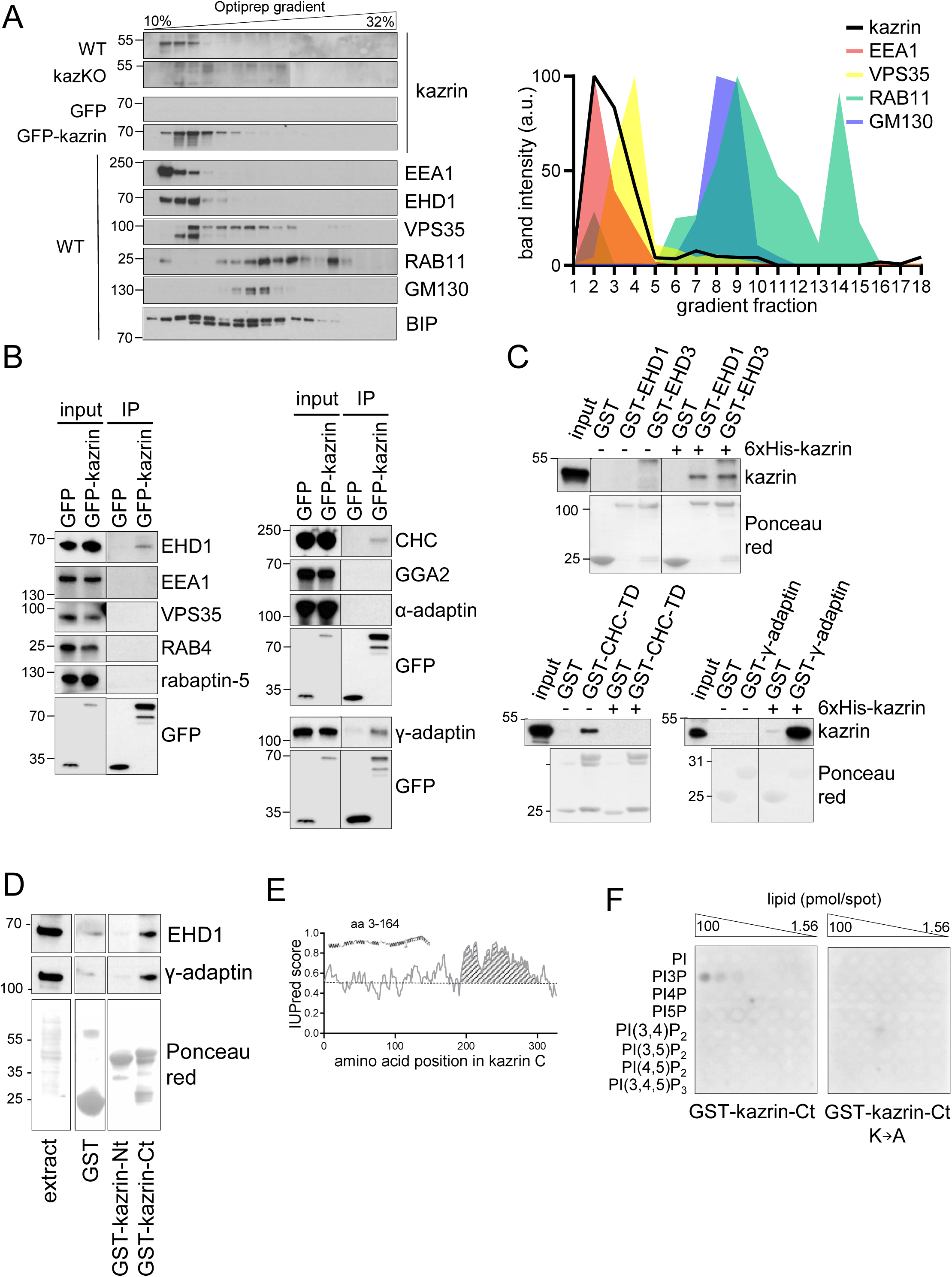
Kazrin is an endosomal protein. (**A**) Left, immunoblots of an Optiprep density gradient fractionation of membrane lysates of WT and kazKO MEF or kazKO MEF moderately (see M & M) expressing GFP or GFP-kazrin C. The membranes were probed with antibodies against the kazrin C N-terminus, EEA1 and EHD1 (EE markers), VPS35 (RAB5/RAB7 transition endosomal marker), RAB11 (RE/Golgi marker), GM130 (*cis*-Golgi marker) and BIP (Binding Immunoglobulin Protein) (ER marker). The antibody against EHD1 is like to recognize other EHD proteins. Band intensity plots per fraction for kazrin or the indicated intracellular membrane markers are shown on the right. The signal intensities of each fraction were normalized to the maximum for each antibody. All gradients were loaded with the same amount of total protein. (**B**) Immunoblots of α-GFP-agarose precipitates from lysates of kazKO MEF moderately expressing GFP or GFP-kazrin C, probed with antibodies against the indicated proteins. 10 µg of total protein were loaded as input. (**C**) Immunoblots of pull-downs from glutathione-Sepharose beads coated with GST, or GST fused to full length EHD1 or EHD3, the Clathrin Heavy Chain terminal domain (CHC-TD) or the γ-adaptin ear domain, incubated with purified 6xHis-kazrin C. The membranes were probed with an α-kazrin antibody (ab74114, from Abcam) and stained with Ponceau red to visualized the GST fusion constructs. (**D**) Immunoblots of pull-downs from glutathione-Sepharose beads coated with GST, or GST fused to the N-(amino acids 1-174) or C-(amino acids 161-327) terminal portions of kazrin C, incubated with non-denaturing extracts from MEFs. 10 µg of total protein were loaded as input. Ponceau red staining of the same membrane (lower panels) is shown to visualize the protein extract or the GST fusion constructs. (**E**) Prediction of IDRs in kazrin C. The graph shows the probability of each residue of being part of an IDR, according to the IUPred2A software (Mészáros, Erdös and Dosztányi, 2018). Residues in the shaded area have a consistent probability over 0.5 to form part of an IDR. (**F**) Immunoblots of a lipid binding assay performed with either the purified GST-kazrin C C-terminal portion (amino acids 161-327) (GFP-kaz-Ct) or an equivalent construct in which the poly-K region has been mutated to poly-A. The membranes used in this assay contain a concentration gradient of the indicated phosphoinositides. Membranes were probed with an α-GST antibody.

To confirm the interaction of kazrin C with endosomes, we immunoprecipitated GFP-kazrin C from native cellular extracts and probed the immunoprecipitates for a number of proteins involved in endosomal trafficking. We detected specific interactions of GFP-kazrin C with EHD proteins, as well as clathrin and γ-adaptin, a component of the Golgi and endosomal clathrin adaptor AP-1 (Adaptor Protein 1) (Fig. 3B). No interaction with the retromer subunit VPS35, the tethering factor EEA1, or the clathrin adaptors GGA2 (Golgi-localised Gamma-ear-containing ADP-ribosylation factor-binding 2) or AP-2 (Adaptor Protein 2) could be demonstrated in immunoprecipitations assays (Fig. 3B), indicating that kazrin C might preferably bind the machinery implicated in endosomal traffic from EEs to or through REs (Grant and Caplan, 2008; Perrin et al., 2013). Pull down assays with purified components indicated that kazrin C can directly interact with EHD1 and EHD3, the clathrin heavy chain terminal domain and the γ-adaptin ear (Fig. 3C). Pull down assays from cell extracts indicated that the EHD proteins and the AP-1 complex bind to the C-terminus of kazrin C, predicted to be an IDR, but not to the N-terminus (Fig. 3D & E). Most interacting partners for kazrin were previously defined to bind its N-terminal region predicted to form a coiled-coil (Groot et al., 2004; Sevilla et al., 2008b). The interaction of endogenous kazrin with γ-adaptin and clathrin could also be confirmed in co-immunoprecipitation experiments from Cos7 and MEFs, using a polyclonal antibody against the C-terminus of kazrin C (Fig. S2C). Also consistent with kazrin specifically interacting with EEs, we found that purified kazrin C interacted with PhosphatidylInositol 3-Phosphate (PI3P) (Fig. 3F), a lipid particularly enriched on EEs (Wang et al., 2019). The interaction required the polyK stretch in the C-terminus of kazrin C (Fig. 3E), previously proposed to constitute a nuclear localization signal (Groot et al., 2004). The data suggested that the C-terminal predicted IDR of kazrin C bears binding sites for multiple EE components, and therefore, it might be required for its EE recruitment and its function in endosomal traffic.

To investigate the role of the C-terminal region of kazrin C in its recruitment to endosomes and its function in endocytic traffic, we generated kazKO cells expressing a GFP-kazrin C construct lacking the C-terminal predicted IDR (lacking amino acids 161 to 327) (kazKO GFP-kazrin C-Nt) using the lentivirus system (Fig. S3A). We then compared its subcellular localization and its capacity to complement the kazKO endocytic defects with those of the full length GFP-kazrin C. As shown in figure 4A, moderately expressed GFP-kazrin C mostly associated with the microsomal fraction containing the EEs, and it was relatively scarce in the cytosol. In contrast, GFP and GFP-kazrin C-Nt appeared more enriched in the cytosolic fraction devoid of membranes (Fig. 4A), indicating that the C-terminal predicted IDR, which binds PI3P, γ-adaptin and EHD proteins, might be required to bring kazrin to cellular membranes. Next, we proceeded to image cells expressing moderate levels of GFP-kazrin C and GFP-kazrin C-Nt, upon loading of the EE with TxR-Tfn at 16°C. The previously reported localization of kazrin C in the nucleus and at cell-cell contacts was evident in these cells (Fig. 4B and S3B; (Groot et al., 2004)). At the PM, GFP-kazrin C neatly co-localized with the *adherens* junction components N-cadherin, β-catenin and p120-catenin, but not with desmoglein, a desmosomal cadherin (Fig. S3B). In addition to the previously reported localizations, GFP-kazrin C concentrated in small condensates, which associated with the surface of the TxR-Tfn labeled endosomes (Fig. 4B & C and movies S3 to S6). Co-localization of GFP-kazrin C condensates with EHD-labeled structures could also be observed in the cell periphery (Fig. S3C and movies S7 to S10). GFP-kazrin C-Nt and GFP staining at similar expression levels appeared mostly cytosolic, with nearly no visible (GFP) or scarce (1 or 2, GFP-kazrin C-Nt) condensates per cell (Fig. 4B & C and movies S11 to S16). The few GFP-kazrin C-Nt condensates observable did not appear associated with TxR-Tn loaded endosomes (Movies S11 to S14).

**Figure 4.**
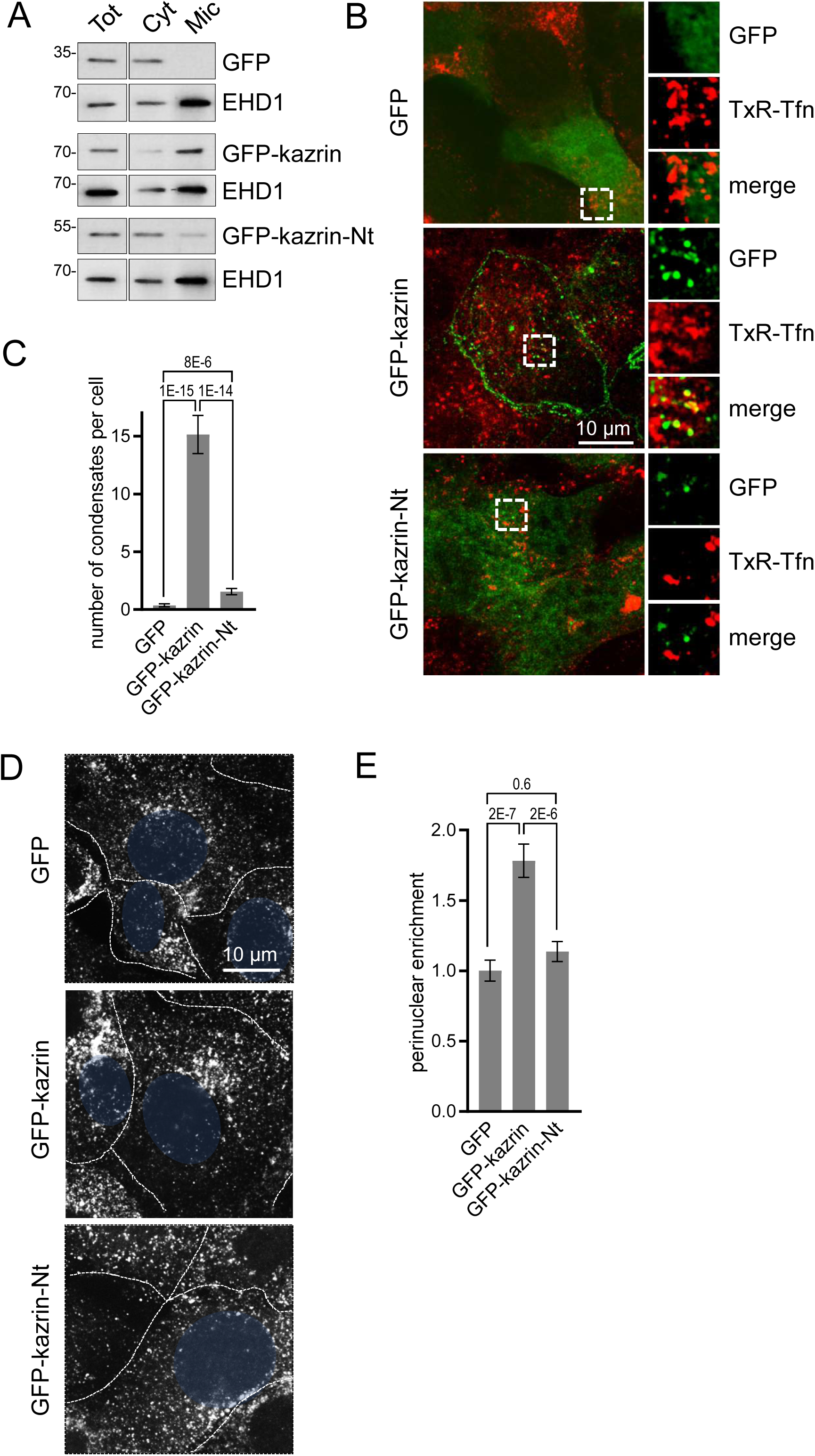
The kazrin C predicted IDR is required for its endocytic function. (**A**) Immunoblots of subcellular fractionations from kazKO cells expressing moderate amounts of GFP, GFP-kazrin C or a GFP-kazrin C construct lacking the C-terminal predicted IDR (GFP-kazrin-Nt). Cells were lysed in a non-denaturing buffer and centrifuged at 186000 g for 1 hour to separate membranes (Mic) from cytosol (Cyt). 15 µg of the supernatant of the 3000 g centrifugation (Tot), and 1 and 5 equivalents of the cytosolic and membrane fractions were loaded per lane, respectively. (**B**) Maximum intensity Z-projections kazKO MEF expressing moderate amounts of GFP, GFP-kazrin C or a GFP-kazrin C construct lacking the C-terminal predicted IDR (GFP-kazrin-Nt), loaded with 20 µg/ml of TxR-Tfn at 16°C to accumulate endocytic cargo on EEs. The images from the GFP and TxR channels and the merge from 5 x 5 µm^2^ fields are shown on the right. (**C**) Mean ± SEM of the number of condensates per cell, visible with the GFP filter channel in the kazKO cells described in (B). p values of two-tailed Mann-Whitney test are shown. n = 29 for each sample. (**D**) Confocal micrographs of kazKO cells expressing low amounts of GFP, GFP-kazrin C or a GFP-kazrin C construct lacking the C-terminal predicted IDR (GFP-kazrin-Nt) loaded with 20 µg/ml of TxR-Tfn at 16°C and chased for 10 min at 37°C. Cell borders are indicated by dashed lines and the nuclei in blue. (**E**) Mean ± SEM TxR-Tfn perinuclear enrichment for the cells and experimental conditions described in D. The mean fluorescence intensity within a circle of 10 µm in the perinuclear region was quantified and divided by the mean signal in the cell. The data is normalized to the mean value of kazKO cells expressing GFP. p values of two-tailed Mann-Whitney test are shown. n > 25 for each sample.

The data thus indicated that the C-terminal predicted IDR was required to recruit kazrin C to endosomal membranes and consequently, it should be required to sustain its function in endosomal traffic, provided that it played a direct role in the process. To test this hypothesis, we investigated the capacity of GFP-kazrin C-Nt to restore the traffic of TxR-Tfn to the perinuclear region in the kazKO cells, as compared to the full length kazrin C. As shown in figures 4D and E, GFP-kazrin C significantly increased the perinuclear enrichment of TxR-Tfn in a kazKO background upon a 10 minute uptake, as compared to GFP expression, whereas GFP-kazrin C-Nt did not.

### Kazrin C localizes at the pericentriolar region and directly interacts with dynactin and LIC1

Interestingly, we observed that in most cells expressing GFP-kazrin C, one or two very bright condensates embracing EE were visible in the perinuclear region (Fig. 5A). Neat co-localization of the bright perinuclear GFP-kazrin C condensates with pericentrin demonstrated that GFP-kazrin C accumulated in the pericentriolar region (Fig. 5B). Live-cell imaging evidenced small GFP-kazrin C condensates moving in and out from the pericentriolar region (Fig. S3D and movie S17). These structures were reminiscent of pericentriolar satellites, which are IDR-enriched membrane-less compartments that transport centrosomal components in a microtubule-dependent manner (Prosser and Pelletier, 2020). Treatment with the microtubule depolymerizing drug nocodazole disrupted the perinuclear localization of GFP-kazrin C, as well as the concomitant perinuclear accumulation of EE (Fig. 5C & D), indicating that EEs and GFP-kazrin C localization at the pericentrosomal region required minus end directed microtubule-dependent transport, mostly effected by the dynactin/dynein complex (Flores-Rodriguez et al., 2011).

**Figure 5.**
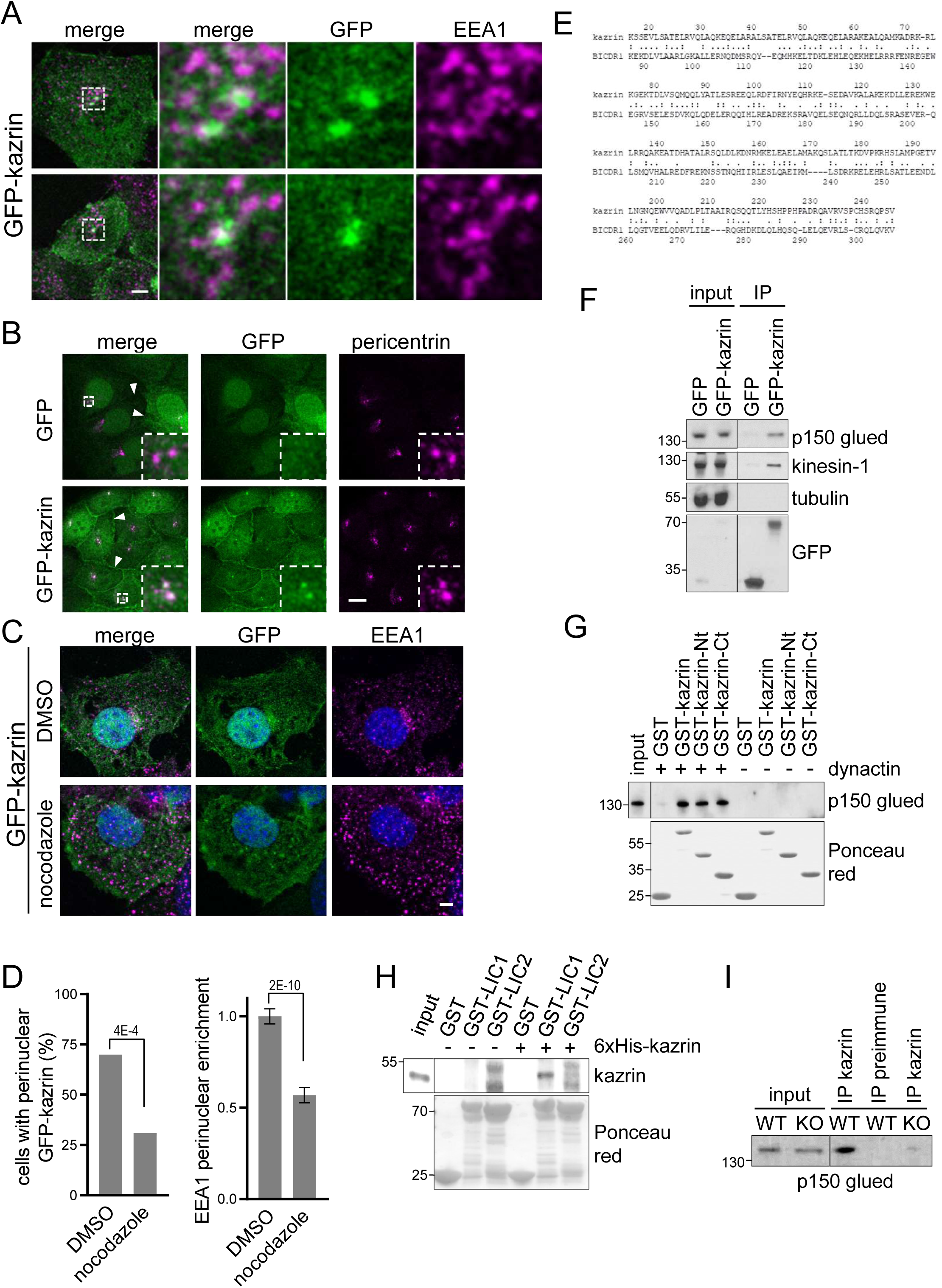
Karin C concentrates in the pericentriolar region and directly interacts with the dynactin complex and the dynein Light Intermediate Chain 1 (LIC1) (**A**) Merged confocal fluorescence micrographs of kazKO MEF moderately expressing GFP-kazrin C, fixed and stained with α-EEA1 and A568-conjugated secondary antibodies. Individual channels and merged confocal images of 6x magnifications are shown. Scale bar, 10 μm. (**B**) Merged and individual channels of confocal fluorescence micrographs of kazKO MEF moderately expressing GFP or GFP-kazrin C, fixed and stained with α-pericentrin and A568-conjugated secondary antibodies. 3.5x magnifications are shown. Arrowheads indicate cell-cell borders. Scale bar, 10 μm. (**C**) Confocal fluorescence micrographs of kazKO MEF moderately expressing GFP-kazrin C, treated with DMSO or 100 ng/ml nocodazole for 16 hours, fixed, and stained with α-EEA1 and A568-conjugated secondary antibodies and DAPI. Scale bar, 5 μm. (**D**) Percentage of cells with a perinuclear localization pattern of GFP-kazrin C (left) and mean ± SEM EEA1 perinuclear enrichment (right) in cells treated as described in C. n > 32 for each sample. The EEA1 perinuclear enrichment was quantified as the mean fluorescence intensity signal within a circle of 10 µm in the perinuclear region divided by the mean cell signal. The data was normalized to the mean of the cells mock treated. The p values of a two-sided Fisher’s exact test (left) and a two-tailed Student t test (right) are shown. n > 32 for each sample. (**E**) Sequence comparison between kazrin C and human BICDR1 calculated with LALIGN. (**F**) Immunoblots of α-GFP agarose immunoprecipitates (IP) from cell lysates of kazKO MEF moderately expressing GFP or GFP-kazrin C, probed with α-p150 glued (dynactin), α-kinesin-1, α-tubulin or α-GFP antibodies. (**G**) Immunoblots of pull-downs with purified GST or GST fused to the full length kazrin C (GST-kazrin) or its N-terminal (amino acids 1-176) (GST-kazrin-Nt) or C-terminal (amino acids 161-327) (GST-kazrin-Ct) portions, incubated with (+) or without (-) the dynactin complex purified from pig. The membranes were probed with a α-p150 glued antibody or stained with Ponceau red to detect the GST constructs. (**H**) Immunoblots of pull-downs with glutathione-Sepharose beads coated with GST, GST-LIC1 or GST-LIC2 incubated with purified 6xHis-kazrin C. The membranes were probed with a mouse α-kazrin antibody or stained with Ponceau red to detect the GST constructs. (**I**) Immunoblot of protein A-Sepharose immunoprecipitates (IP) from WT or kazKO MEFs using a mix of rabbit polyclonal serums against the N- and C-terminal domains of kazrin C or a pre-immunisation serum, probed with a α-p150 glued (dynactin) antibody. The low amounts of endogenous kazrin could not be detected in the immunoprecipitates with any of the antibodies tested because the antibody had a molecular weight similar to endogenous kazrin (about 50 Kda) and interfered with the detection. The kazKO MEF were used as a specificity control instead.

Our observations indicated that kazrin C can be transported in and out of the pericentriolar region along microtubule tracks, and that it is required for the perinuclear accumulation of EEs. Interestingly, pericentriolar localization of GFP-kazrin C was reminiscent of that observed for well-established or candidate dynein/dynactin activating adaptors such as hook2 or hook3 (Baron and Salisbury, 1988; Szebenyi et al., 2007). Indeed, kazrin C shared 23.3% identity and 57.3 % similarity with BICDR1 (BICauDal Related protein 1) (Fig. 5E), over 232 amino acids, and 19.6% identity and 61.3 % similarity with hook3, over 168 amino acids. Such similarity was in the range of that shared between hook3 and BICDR1 (24.7% identity and 56.7 % similarity over 268 amino acids) (LALIGN). Therefore, we hypothesized that kazrin might also interact with the dynein/dynactin complex and serve as an endosomal dynein adaptor. Consistent with this hypothesis, we found that moderately expressed GFP-kazrin C pulled-down the dynactin component p150-glued from cell extracts, whereas GFP alone did not (Fig. 5F). Similar to what has been described for other dynein/dynactin adaptors (Kendrick et al., 2019), we also detected co-immunoprecipitation of GFP-kazrin C with plus-end directed motors, specifically, with kinesin-1 (Fig. 5F), a motor associated with EEs (Loubery et al., 2008). We observed no co-immunoprecipitation with tubulin (Fig. 5F), indicating that GFP-kazrin C interactions with dynactin and kinesin-1 were not indirectly mediated by microtubules. Pull down experiments with GST-kazrin C, expressed and purified from *E. coli,* and the dynactin complex, purified from pig (Jha et al., 2017), demonstrated that the interaction was direct and that it was contributed by both, the N- and C-terminal halves of kazrin C (Fig. 5G), suggesting multiple contacts with different dynactin components. As also described for other dynein/dynactin adaptors such as BICD2, CRACR2a (Calcium Release Activated Channel Regulator 2a) and hook3 (Lee et al., 2020), pull down experiments with purified components evidenced a weak, albeit specific interaction of kazrin C with the dynein LIC1, but not with LIC2 (Fig. 5H). Finally, immunoprecipitation experiments from MEFs using the polyclonal antibody against the C-terminus of kazrin C also suggested binding of endogenous kazrin with the dynactin complex (Fig. 5I).

Our data indicated that kazrin might act as a new endosomal dynein/dynactin adaptor, with its C-terminal IDR working as a scaffold that entraps EE or certain EE subdomains through multiple low affinity binding sites. To directly test this hypothesis we applied high speed live-cell fluorescence imaging to visualize the movement of TxR-Tfn-loaded EEs in WT and kazKO cells. As previously described, EEs in WT cells were highly motile exhibiting long range trajectories of several micrometers, followed by more confined movements within a 1 µm radius (Flores-Rodriguez et al., 2011; Loubery et al., 2008) (Movie S18). Kymographs of maximum intensity Z-stack projections of 90 seconds movies evidenced the linear endosomal trajectories in WT cells, with an average length of about 5 µm (Fig. 6A & B). However, in kazKO MEFs, the kymographs showed profusion of bright dots as compared to the straight trajectories in the WT, and the length of the straight trajectories (longer than 1 µm) was significantly reduced (Fig. 6A & B and movies S18 & S19). These observations suggested that the absence of kazrin reduced the association of EEs with some microtubule-dependent motors and/or diminished the processitiy of those (Fig. 6A & B). Analysis of the maximum instantaneous velocities (Vi) of centripetal trajectories longer than 1 µm, mostly contributed by dynein (Flores-Rodriguez et al., 2011; Loubery et al., 2008), showed that those were lower in the kazKO cells, as compared to the WT (Fig. 6C & movies S18 to S19). Finally, and also supporting the view that kazrin directly contributes to EE centripetal transport, we observed that expression of GFP-kazrin C, but not expression of the truncated version lacking the C-terminal endosomal binding region (GFP-kazrin C-Nt) nor GFP alone, rescued the endosome motility defects installed by depletion of kazrin (Fig. 6A to C, and movies S21 to S22).

**Figure 6.**
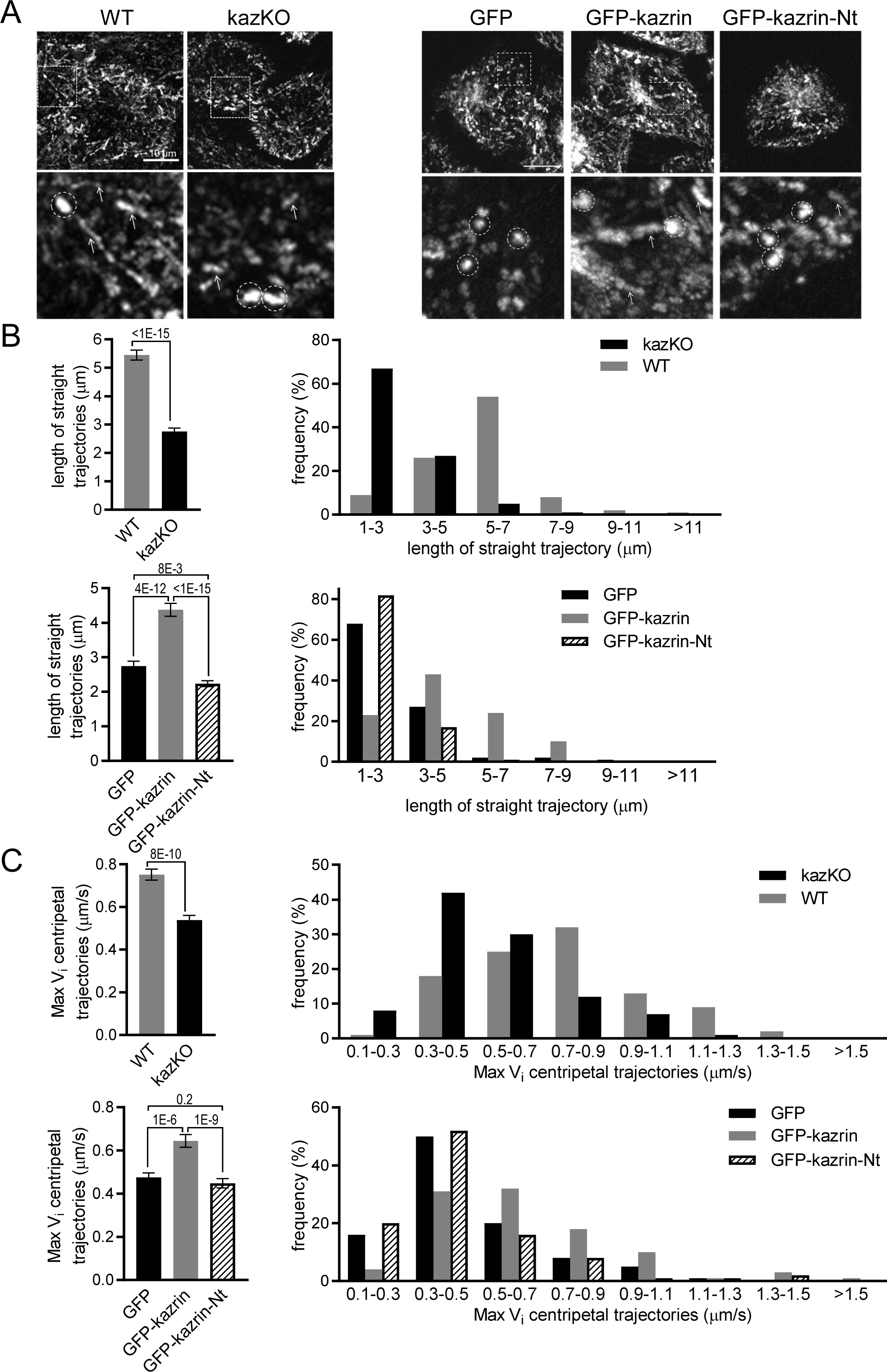
Depletion of kazrin impairs endosome motility. (**A**) Kymographs of maximum intensity Z projections of confocal fluorescence microscopy movies taken for 90 seconds with a 3 second time lapse, of WT and kazKO MEF or kazKO MEF expressing low levels of GFP, GFP-kazrin C of a GFP-kazrin C construct lacking the C-terminal predicted IDR (GFP-kazrin-Nt), showing trajectories of EE loaded with 20 µg/ml of TxR-Tfn at 16℃. Cells were shifted to 37℃ and immediately imaged. A magnified 10 x 10 µm^2^ inset is shown below. Arrows point to straight trajectories and dashed circles indicate constrained endosome movements. (**B**) Mean ± SEM (left graphs) of the length of straight endosome trajectories (longer than 1µm) for the cells and experimental conditions described in (A). p values of two-tailed Mann-Whitney tests are shown. n = 100 for each sample. Frequencies of the trajectory length in each cell type are shown on the right. (**C**). Mean ± SEM (left graphs) of the maximum instantaneous velocities (Vi) of centripetal endosome trajectories (longer than 1µm) for the cells and experimental conditions described in (A). p values of two-tailed Mann-Whitney tests are shown. n = 100 for each sample. Frequencies of the maximum Vi for each cell type are shown on the right.

## Discussion

Our data suggests the kazrin plays a primary role in endosomal recycling through the long pathway traversing the perinuclear region, and that it does so by working as a dynein/dynactin adaptor for EEs or EE-derived transport intermediates. We showed that the predicted kazrin IDR directly interacts with several EE components implicated in endocytic traffic (Grant and Caplan, 2008; Perrin et al., 2013) and that depletion of kazrin causes a defect in the transport of endocytosed Tfn to the perinuclear region, as well as the scattering of EEs but not REs to the cell periphery. These phenotypes recapitulate those observed when depleting other proteins involved in EE to RE transport such as EHD3, or upon inhibition of dynein (Driskell et al., 2007; Naslavsky et al., 2006; Nielsen et al., 1999). We also showed that kazrin shares considerable homology to dynein/dynactin adaptors and that its depletion or deletion of its IDR impairs movement of Tfn-loaded endosomes towards the perinuclear region.

While the vesicular nature of membrane traffic from the PM to the EE has been well characterized, the principles governing the transport of membranes and cargo within the endosomal system are much less understood. The more accepted view is that the core of the EEs, receiving the internalized material, undergoes a maturation process that leads to its conversion to LEs, while retrograde traffic is driven by tubular transport intermediates, generated by sortinexins (SNX) or clathrin coated vesicles (Haberg et al., 2008; Hsu et al., 2012; McNally and Cullen, 2018). In addition, centripetal transport of EEs to the pericentriolar region has been proposed to facilitate fusion with or maturation to REs (Naslavsky and Caplan, 2018; Solinger et al., 2020).

Within this wide range of endosomal trafficking events, microtubules seem to play key roles. EEs move along microtubule tracks with a bias toward the cell center (Driskell et al., 2007; Nielsen et al., 1999). Centripetal movement is mainly effected by the dynein/dynactin complex (Granger et al., 2014). Treatment of cells with nocodazol, or interfering with dynein, results in inhibition of endosome motility, the scattering of EEs to the cell periphery and impaired endosomal maturation (Granger et al., 2014). In addition, plus and minus-end directed microtubule-dependent-motors have both a role in maintaining the endosomal subdomain organization and in the formation and motility of SNX-dependent tubular structures (Granger et al., 2014). Interestingly, motility of different SNX tubules or endosomal subdomains is associated with distinct dynein complexes bearing either the LIC1 or LIC2 chains and particular kinesin types (Hunt et al., 2013). In this context, the interactome of kazrin C suggests that it might work as a LIC1-dynein and kinesin-1 adaptor for EHD and/or AP-1/clathrin transport intermediates emanating from EEs, in transit to the RE (Grant and Caplan, 2008; Perrin et al., 2013). Hook1 and Hook3, as components of FHF (Fused Toes-Hook-Fused toes and Hook Interacting Protein) complexes, have also been proposed to work as EEs dynein/dynactin adaptors in yeast, fruit flies and mammalian cells ((Olenick and Holzbaur, 2019; Xiang and Qiu, 2020; Xiang et al., 2015) and references therein). However, the interactome of the mammalian hook1 and hook3 and the endocytic pathways affected by interfering with their function differ from those of kazrin (Christensen et al., 2021; Guo et al., 2016; Maldonado-Baez and Donaldson, 2013; Olenick et al., 2019; Xu et al., 2008).

The role of kazrin in endocytic recycling might explain some of the pleiotropic effects observed in vertebrate development upon its depletion. Defects in the establishment of cell-cell contacts in *Xenopus* embryos and human keratynocytes (Cho et al., 2010; Sevilla et al., 2008a; Sevilla et al., 2008b) might derive from altered recycling of cadherins or desmosomal components (Kawauchi, 2012). Indeed, depletion of kazrin in *Xenopus laevis* leads to decreased levels of E-cadherin, which can be reverted by inhibiting endocytic uptake (Cho et al., 2010). This observation is consistent with a role of kazrin diverting traffic of internalized E-cadherin away from the lysosomal compartment and back to the PM. Likewise, eye and craniofacial defects associated with reduced EMT and neural crest cell migration (Cho et al., 2011), might originate from altered endocytic trafficking of integrins, cadherins and/or signaling receptors (Cadwell et al., 2016; Jones et al., 2006; Wilson et al., 2018).

It is worth noticing that kazrin is only expressed in vertebrates, whose evolution is linked to an explosion in the number of cadherin genes and the appearance of desmosomes (Green et al., 2020; Gul et al., 2017). In this context, it is tempting to speculate that while the core machinery involved in membrane traffic is largely conserved from yeast to humans, vertebrates might have had the need to develop specialized trafficking machinery such as kazrin, which spatiotemporally regulates the function of particular adhesion complexes. Therefore, kazrin might turn out to be a valid therapeutic target to selectively modulate the function of those adhesion complexes in the context of a myriad of human pathologies (Kaszak et al., 2020; Yuan and Arikkath, 2017). Identification of the relevant endocytic cargo travelling in a kazrin-dependent manner will be the next step to further understand the molecular, cellular and developmental functions of kazrin.

## MATERIALS AND METHODS

### DNA techniques and plasmid construction

Oligonucleotides used for plasmids construction and information about the construction strategies are available upon request. DNA manipulations were performed as described (Sambrook et al., 1989), or with the Getaway cloning system (Life Technologies) in the case of lentiviral vectors. Enzymes for molecular biology were obtained from New England Biolabs. Plasmids were purified with the Nucleospin plasmid purification kit (Macherey-Nagel). Linear DNA was purified from agarose gels using the gel extraction kit from Qiagen. Polymerase Chain Reactions (PCRs) were performed with the Expand High Fidelity polymerase (Roche) and a TRIO-thermoblock (Biometra GmbH). Plasmids used are listed in Table I.

**Table I.**
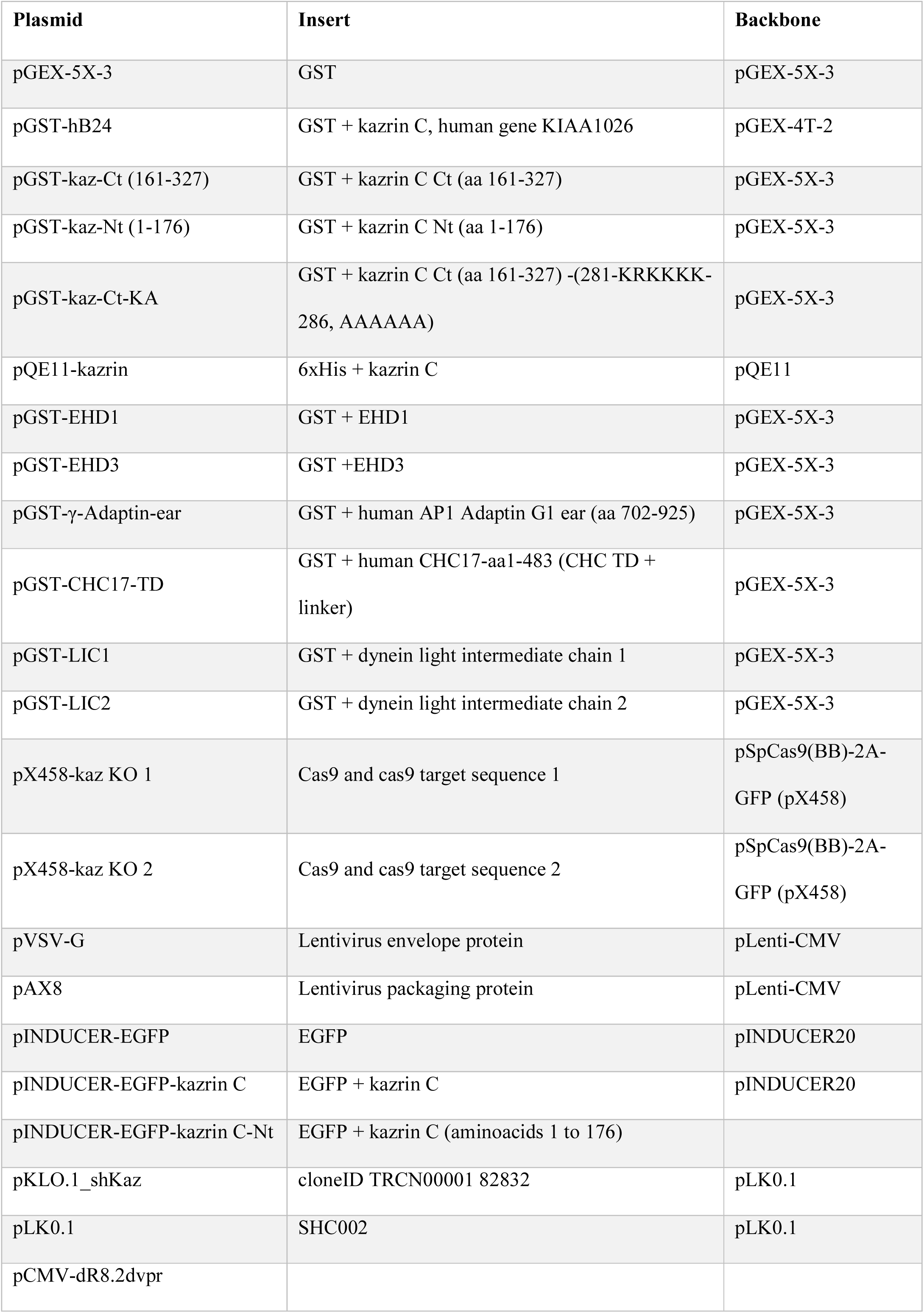
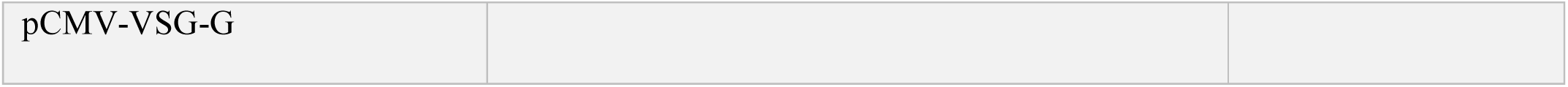
Plasmids.

| **Plasmid** | **Insert** | **Backbone** |
| --- | --- | --- |
| pGEX-5X-3 | GST | pGEX-5X-3 |
| pGST-hB24 | GST + kazrin C, human gene KIAA1026 | pGEX-4T-2 |
| pGST-kaz-Ct (161-327) | GST + kazrin C Ct (aa 161-327) | pGEX-5X-3 |
| pGST-kaz-Nt (1-176) | GST + kazrin C Nt (aa 1-176) | pGEX-5X-3 |
| pGST-kaz-Ct-KA | GST + kazrin C Ct (aa 161-327) -(281-KRKKKK-286, AAAAAA) | pGEX-5X-3 |
| pQE11-kazrin | 6xHis + kazrin C | pQE11 |
| pGST-EHD1 | GST + EHD1 | pGEX-5X-3 |
| pGST-EHD3 | GST + EHD3 | pGEX-5X-3 |
| pGST-γ-Adaptin-ear | GST + human AP1 Adaptin G1 ear (aa 702-925) | pGEX-5X-3 |
| pGST-CHC17-TD | GST + human CHC17-aa1-483 (CHC TD + linker) | pGEX-5X-3 |
| pGST-LIC1 | GST + dynein light intermediate chain 1 | pGEX-5X-3 |
| pGST-LIC2 | GST + dynein light intermediate chain 2 | pGEX-5X-3 |
| pX458-kaz KO 1 | Cas9 and cas9 target sequence 1 | pSpCas9(BB)-2A-GFP (pX458) |
| pX458-kaz KO 2 | Cas9 and cas9 target sequence 2 | pSpCas9(BB)-2A-GFP (pX458) |
| pVSV-G | Lentivirus envelope protein | pLenti-CMV |
| pAX8 | Lentivirus packaging protein | pLenti-CMV |
| pINDUCER-EGFP | EGFP | pINDUCER20 |
| pINDUCER-EGFP-kazrin C | EGFP + kazrin C | pINDUCER20 |
| pINDUCER-EGFP-kazrin C-Nt | EGFP + kazrin C (aminoacids 1 to 176) |  |
| pKLO.1_shKaz | cloneID TRCN00001 82832 | pLK0.1 |
| pLK0.1 | SHC002 | pLK0.1 |
| pCMV-dR8.2dvpr |  |  |
| pCMV-VSG-G |  |  |

### Cell culture and cell line establishment

Cos7 cells were obtained from the German Collection of Microorganisms and Cell Cultures (https://www.dsmz.de/dsmz). MEF and HEK293T cells were provided by A. Aragay (IBMB-CSIC). Cells were grown in DMEM (Thermo Fisher) supplemented with 10 % FBS, 100 µ/ml penicillin, 100 µg/ml streptomycin and 2 mM L-glutamine (Thermo Fisher) in a humidified 5 % CO_2_ atmosphere at 37°C. Cos7 were transiently transfected with Lipofectamine 2000 (Thermo Fisher). Cells were analyzed 24 hours after transfection. For shRNA kazrin depletion, pKLO.1_shKaz from Merck Mission Library 2007 (Clone ID TRCN000018283) was used. pLKO.1_CV/_SCR (SHC002) was used as a control. For lentivirus production and Cos7 cells transfection, HEK293T cells were co-transfected with either the pLL3.7 encoding GFP, for virus production control and infection efficiency monitoring, or with pLKO.1 encoding the desired shRNA, and the viral packaging (pCMV-dR8.2 dvpr) and envelope (pCMV-VSV-G) plasmids, using calcium phosphate transfection. About 16 hours after transfection, the medium was changed and half of the usual volume was added. During the two following days, medium containing the virus was collected and filtered with a 0.45 μm filter (Millipore). The filtered virus solution was directly used for the infection of cell lines or stored in aliquots at −80°C without prior concentration of the virus. Infection and selection of stably infected cells were done in the presence of the appropriate concentration of puromycin (Merck-Aldrich), titrated by using the minimum antibiotic concentration sufficient to kill untransfected cells, but to maintain cells transfected with the pLL3.7 GFP-containing plasmid. Actual depletion of kazrin or the protein of interest was analyzed by immunoblot using home-made polyclonal rabbit antibodies raised against the N- (amino acids 1 to 176) and C- terminal (amino acids 161 to 327) portions of kazrin C.

MEF KO cells were produced with the CRISPR-CAS9 system. Two guide RNAs were designed to recognize regions at the start of exon 2 of the kazrin gene, corresponding to the start of kazrin C isoform (CACCGAATGCTGGCGAAGGACCTGG and CACCGCCTTCTGTACCAGCTGCACC). Online tools Benchling (https://www.benchling.com/) and the Broad Institute tool GPP (https://portals.broadinstitute.org/gpp/public/analysis-tools/sgrna-design) were used for the design. Guide RNA oligonucleotides were annealed and inserted into a pSpCas9(BB)-2A-GFP pX458 vector. MEFs were electroporated with Nucleofector (Lonza), following the manufacturer’s instructions. GFP-positive cells were sorted by FACS in an Aria FUSION (Becton Dikinson) sorter and screened by immunoblotting with antibodies against the N-terminal and the C-terminal portions of kazrin. Lentiviral particles were produced in HEK 293T cells. Calcium phosphate-mediated transfection was used to deliver vector pINDUCER20 encoding GFP or GFP-tagged kazrin constructs, together with packaging and envelope lentiviral vectors. The supernatant of transfected HEK 293T cells was collected after 16 hours, 0.45 µm-filtered and added to MEFs. The cells were passaged for a week, incubated with 5 µg/ml doxycycline (Millipore) for 48 hours to induce the expression of the construct. GFP-positive cells were selected by FACS and pooled. MEFs were transfected by electroporation using the Ingenio solution (Mirus) and a nucleofector (Amaxa). The cells were processed 2 days after electroporation.

For complementation assays, GFP, GFP-kazrin C and GFP-kazrin C-Nt were induced for up to 12 hours to achieve low, nearly-physiological expression levels of GFP-kazrin C (as compared to endogenous kazrin by immunoblot, using the home-made rabbit polyclonal anti-kazrin serums), and analogous expression levels of GFP or GFP-kazrin C-Nt (as compared by immunoblot using the mouse anti-GFP antibody (see antibodies section)). For GFP-kazrin C imaging or biochemical studies, cells were induced for up to 24 hours to achieve analogous, moderately-overexpressed levels of the proteins. To study the effect on microtubule dynamics, MEFs were treated with 100 ng/ml of nocodazole (Merck) or DMSO for 16 h, and then fixed at room temperature.

### TxR-Tfn accumulation, perinuclear enrichment, and recycling assays

Cos7 cells or MEFs were grown on R-collagen-coated glass coverslips. For all assays, cells were starved 30 minutes in DMEM without FBS or BSA. For the accumulation assays, cells were then incubated with pre-warmed DMEM containing 20 µg/ml of TxR-Tfn (from human serum, Molecular Probes) and 0.1 % BSA for the specified times. Cells were washed in ice-cold PBS once and fixed in 4 % PFA (Merck) for 20 minutes on ice. For the TxR-Tfn perinuclear enrichment and recycling assays, 20 µg/ml of TxR-Tfn in DMEM with 0.1 % BSA was added and cells were incubated at 16 °C for 30 minutes to load EEs. Cells were then washed in ice-cold PBS with 25 mM acetic acid pH 4.2, and with PBS and subsequently incubated with 500 µg/ml unlabeled Tfn (Merck) in DMEM with 0.1 % BSA at 37 °C. Cells were then transferred to ice at the indicated time points, washed in ice-cold PBS with 25 mM acetic acid pH 4.2 and with PBS, and fixed in 4 % PFA for 20 minutes on ice. For the perinuclear enrichment assays the mean TxR-Tfn fluorescence intensity within a 10 µm diameter circle in the perinuclear region was divided by the signal in the whole cell selected with the Fiji free hand tool to define the ROI (Regio n Of Interest), at 10 minutes chase, after background subtraction. For recycling experiments the mean fluorescence intensity per cell was measured using the Fiji free hand drawing tool to select the ROI at the indicated time points and the signal was normalized to the average intensity at time 0.

For TxR-Tfn accumulation, perinuclear enrichment and recycling assays, images were taken with a Zeiss LSM780 confocal microscope equipped with a 63x oil (NA = 1.4) objective, a GaAsP PMT detector 45% QE and images were acquired at pixel size 0.06 µm, unless otherwise indicated. For the experiments shown in figure 4D, an Andor Dragonfly spinning disk microscope equipped with a 100x oil (NA = 1.49) objective and a Sona 4.2 B11 sCMOS camera 95% QE was used. Images were acquired at pixel size 0.05 µm.

### Colocalization of GFP-kazrin and TxR-Tfn and immunofluorescence

3D reconstructions of EEs loaded with TxR-Tfn in cells expressing GFP, GFP-kazrin C or the GFP-kazrin C-Nt were performed with voxel size 0.05 x 0.05 x 0.10 µm, compiled with the Andor Dragonfly spinning disk microscope equipped with a 100x oil (NA = 1.49) objective and a Sona 4.2 B11 sCMOS camera 95% QE, in cells treated as for the TxR-Tfn recycling assay, immediately upon the shift from 16℃ to 37℃. 3D movies of 5 x 5 µm^2^ were generated with the Fiji 3D reconstruction tool. A 1.0 Gaussian blur filter was applied to the images after performing the 3D reconstruction. For immunofluorescence experiments, cells were seeded onto cover-glasses and fixed with 4 % PFA in PBS containing 0.02 % BSA and 0.02 % sodium azide (PBS*), for 10 minutes at room temperature. Cells were washed 3 times for 5 minutes with PBS* and permeabilized with PBS* containing 0.25 % Triton X-100 for 10 min. Cells were washed 3 times for 5 minutes with PBS* and incubated for 20 minutes in PBS* containing 1 % BSA. Cells were then incubated in the presence of the primary antibody in PBS* for 1 hour at room temperature, washed 3 times with PBS* and incubated for 1 hour in the presence of the secondary antibodies prepared in PBS*. Cells were washed 3 times with PBS* and mounted using Prolong Gold that included DAPI for nuclear staining (Thermo Fisher). Images were taken with a Zeiss LSM780 confocal microscope equipped with a 63x oil (NA = 1.4) objective, a GaAsP PMT detector 45% QE, and images were acquired at pixel size 0.06 µm for the experiments shown in figures 5B and C and 0.120 µm for the experiments shown in figure 5A. Images shown in figure S3B and the associated movies for the 3D reconstruction of EHD labeled endosomes, were performed with the Andor Dragonfly spinning disk microscope equipped with a 100x oil (NA = 1.49) objective and a Sona 4.2 B11 sCMOS camera 95% QE, with voxel size 0.05 x 0.05 x 0.10 µm. Experiments shown in figure S3B were acquired with a Leica TCS-SP5 confocal microscope equipped with a 63x oil objective (NA = 1.4), with a pixel size of 0.06 µm. Perinuclear enrichments for EEA1 and RAB11 in MEFs were calculated after background substraction as the mean fluorescence intensity within a 10 and 9 µm (respectively) diameter circle in the perinuclear region, divided by the mean intensity in the whole cell, as delimited with the Fiji free hand drawing tool to select the ROI.

### Cell migration and division assays

Cells were plated on 400 µg/ml Matrigel (Corning)-coated plates at low density and incubated for 5 hours. Once the cells were attached, the medium was replaced by Matrigel for 30 minutes to embed the cells in a matrix. Matrigel excess was then removed and cells were kept at 37 °C with 5 % CO_2_ during imaging. Phase contrast images were taken every 10 minutes for a total of 9 hours with a monotorized bright field Leica AF7000 microscope equipped with a 10x objective (NA = 0.3), and a digital Hamamatzu ORCA-R2 CCD camera and images were taken with a pixel size of 0.64 µm. To analyze cell migration, cells were tracked using the Fiji plugin MTrackJ. Speed and direction persistency was calculated using the open-source program DiPer (Dang et al., 2013). To detect cytokinesis delay compatible with a defect in abscission, the time was measured from the moment daughter cells attach to the substrate until they completely detach from each other.

### Live confocal imaging

Cells were seeded on plates with polymer coverslips for high-end microscopy (Ibidi). Cells were kept at 37°C with 5 % CO_2_ during the imaging. For the movie S17 and the figure S3C, images were taken every 2.65 seconds on a Zeiss LSM780 confocal microscope equipped with a 63x oil objective (NA = 1.4) for, with voxel size 0.05 x 0.05 x 0.130 µm. To follow EE motility, cells were starved for 30 minutes in DMEM without FBS and subsequently loaded at 16°C with 20 µg/ml TxR-Tfn in DMEM with 0.1% BSA, as described for the TxR-Tfn recycling experiments. Cells were then rinsed with PBS and image immediately upon addition of 37℃ pre-warmed media loaded with unlabeled Tfn. Images were compiled with voxel size 0.17 x 0.17 x 0.46 µm for WT and KO cells and 0.09 x 0.9 x 0.46 µm for GFP GFP-kazrin C and GFP-kazrin C-Nt expressing cells, and they were taken every 3 seconds for 1.5 minutes using the Andor Dragonfly 505 microscope, equipped with a 60x oil (NA = 1.4) objective and a Sona 4.2 B11 sCMOS camera 95% QE. Maximum intensity projections of the Z-stacks were generated with Fiji, after background subtraction and registration using the Linear Stack Alignment with SIFT tool of Fiji. Movies of 10 x 10 µm^2^ were generated from the original movies using the Fiji crop tool and a 1.0 Gaussian filter was applied. Kymographs of the maximum intensity Z-stack projections were generated to measure the length of linear trajectories with the Fiji free hand line tool. Maximum instantaneous velocity (Vi) of TxR-Tfn loaded endosomes was measured by manually tracking endosomes moving into the cell center with the Fiji plugin MTrackJ.

### SDS-PAGE and immunoblots

SDS–PAGE was performed as described (Laemmli, 1970), using pre-casted Mini-PROTEAN TGX 4-20% Acrylamide gels (Bio Rad). Protein transfer, blotting and chemiluminescence detection were performed using standard procedures. Detection of proteins was performed using the ECL kit (GE Healthcare).

### Cell fractionation

Cell fractionation was performed as described in Li and Donowitz (2014) (Li and Donowitz, 2014). Briefly, cells were scraped from the plate, harvested by centrifugation at 700 g for 10 minutes and resuspended in 1 ml of ice cold Lysis Buffer (LB: 25 mM Hepes pH 7.4, 150 mM NaCl, 1 mM DTT, 2 mM EGTA) containing protease inhibitors. The cell suspension was then passed 10 times through a 27 G needle. The lysate was cleared by centrifuging twice at 3000 g for 15 min. The supernatant was subsequently centrifuged at 186000 g for 1 hour at 4°C to fractionate cellular membranes from cytosol. The membrane pellet was resuspended in LB with protease inhibitors, passed 10 times though a 27 G needle and laid on an Optiprep (Merck) gradient. A 12 ml 2 % step Optiprep gradient in LB ranging from 32 % to 10 % was prepared beforehand in Ultra-Clear tubes (Beckman Coulter). Samples were spun for 16 hours at 100000 g at 4 °C. 0.6 ml fractions were carefully collected from the top. Samples were then precipitated with trichloroacetic acid, air-dried and resuspended in SDS-PAGE sample buffer for immunoblot analysis. For the experiments shown in figure 4A, the supernatant from the 3000 g centrifugation was adjusted to 1 mg/ml of total protein and centrifuged at 186000 g for 1 hour at 4°C to fractionate cellular membranes (pellet) from cytosol (supernatant). 15 µg of total protein from the 3000 g supernatant (total) and the corresponding 1 and 5 equivalents of the cytosolic or membrane fractions were loaded in an SDS-PAGE acrylamide gel and immunoblotted for EHD proteins or GFP.

### GST pull downs, GFP-trap and endogenous immunoprecipitations

Purification of recombinant GST and 6-His fusion proteins from BL21 *E. coli* (Novagen) was performed as described (Geli et al., 2000). Pull down experiments were performed with Glutathione-Sepharose beads (GE Healthcare) coated with 0.5 μg of the indicated GST-tagged proteins and 2 nM of eluted 6xHis-kazrin C incubated in 1 ml of binding buffer containing PBS or 2 nM of the dynactin complex in 0.5 ml of DBB (25 mM Tris-HCl pH 8, 50 mM KoAc, 0.5 mM ATP, 1 mM DTT, 1 mM MgCl_2_, 1 mM EGTA and 10 % glycerol), both bearing 0.2 % Triton-X100 and 0.5 % BSA with protease inhibitors (Complete Roche), for 1 hour at 4 °C in a head-over-shoulder rotation. Beads were washed 3 times with the corresponding binding buffer containing Triton-X100 and twice without detergent. The beads were boiled in Laemmli buffer. Input and pulled-down samples were loaded in an SDS-PAGE gel and analyzed by immunoblot. For the pull downs from mammalian protein extracts, GST, and the GST-kazrin C N- (amino acids 1 to 176) and C-terminal (amino acids 161 to 327) portions were expressed and purified from *E. coli* as described above, using glutathione-Sepharose beads, and the beads were incubated with the 3000 g supernatant of a non-denaturing protein extract from WT MEF, prepared as described for the subcellular fractionation using LB, after adding 1% Triton-X100. After 1 hour incubation, beads were recovered and washed with LB 1% Triton-X100 3 times and twice with LB buffer. Beads were resuspended in SDS-PAGE sample buffer and analyzed by immunoblot against EHD proteins and γ-adaptin.

For immunoprecipitations from MEFs, moderately overexpressing GFP and GFP-kazrin C, the cells were harvested and cleared at 100 g. The pellet was resuspended in 500 µl of IP buffer (20 mM Hepes, 50 mM KAc, 2 mM EDTA, 200 mM sorbitol, 0.1 % Triton X-100, pH 7.2) containing protease inhibitors and passed 30 times through a 27 G needle. The lysate was cleared by centrifuging 5 minutes at 10000 g. 10 µl of GFP-binding agarose beads (Chromotek) were incubated with the protein extract for 1 hour at 4 °C in head-over-shoulder rotation. Beads were washed six times with 1 ml of IP buffer. The beads were boiled in Laemmli buffer. Input and IP samples were loaded in an SDS-PAGE gel and analyzed by immunoblot. DBB (25 mM Tris-HCl pH 8, 50 mM KoAc, 0.5 mM ATP, 1 mM DTT, 1 mM MgCl_2_, 1 mM EGTA and 10 % glycerol) containing 0.1 % Triton X-100 was used for the immunoprecipitation experiments with dynactin and kinesin-1.

For endogenous immunoprecipitations, WT or kazKO cell extracts were generated as described above but incubated with rabbit IgGs against the kazrin C C-terminus (aa 161 to 327), pre-bound from a serum to Protein A-Sepharose, or IgGs from the pre-immune serum. The amount of endogenous kazrin in the immunoprecipitates could not be assessed because the IgGs interfered with the detection.

### Lipid strip assays

Lipid strips (Echelon) were incubated in 1 % skimmed milk in PBS for 1 hour at room temperature. The corresponding GST fusion protein was added to a final concentration of 15 µg/ml in incubation buffer (10 mM Tris pH 8.0, 150 mM NaCl, 0.1 % Tween-20, 3 % BSA (fatty acid free, Merck)), with protease inhibitors over night at 4°C. The strips were washed three times for 10 minutes in the incubation buffer and developed by immunoblot.

### Quantification, statistical analysis and structure prediction

Quantifications were performed with the Fiji open source platform (Schindelin et al., 2012). Statistical analysis was performed with GraphPad Prism. The D’Agostino-Pearson test was applied to data sets to assess normality. If the data followed a normal distribution or the result of the normality test was not significant, an unpaired two-tailed Student t test was performed to assess significance. If the distribution was not normal, a two-tailed Mann-Whitney test was used. Results are expressed as mean ± SEM with respect to the number of cells (n) for a representative experiment. Prediction of IDRs was achieved with the IUPred2A software, which assigns each residue a IUPred score that is the probability of it being part of a IDR (Mészáros, Erdös and Dosztányi, 2018).

### Antibodies

Polyclonal sera against kazrin for immunoblotting were generated in rabbit using an N-terminal (amino acids 1 to 176) and a C-terminal (amino acids 161 to 327) fragment of kazrin C fused to GST. The following commercial antibodies were used in this study: anti-RAB11 (610656), anti-RAB4 (610888), anti-rabaptin-5 (610676), anti-GM130 (610822), anti-GGA2 (612612), anti-clathrin heavy chain (610499), anti-p150 Glued (610473), anti-α-adaptin (610501), anti-γ-adaptin (610386), anti-N-cadherin (51-9001943), anti-β-catenin (610153), anti-p120-catenin (51-9002040) and anti-desmoglein (51-9001952) from BD Biosciences, anti-pericentrin (4448) and anti-EHD1 (109311) from Abcam, anti-VPS35 (374382) and anti-kinesin-1 heavy chain (133184) from Santa Cruz Biotechnology, anti-EEA1 (3288) from Cell Signalling Technology, anti-tubulin (T-6557) from Merck, anti kazrin C (ab74114) from Abcam, anti-GFP (632380) from Living Colors and anti-GST (27-57701) from Amersham. Peroxidase-conjugated anti-mouse (A2554), anti-goat (A4174) and rabbit (A0545) IgGs were from Merck. Alexa Fluor 568 anti-mouse IgG (A11037), Alexa Fluor 568 anti-rabbit IgG (A11036) and Alexa Fluor 647 anti-rabbit IgG (A21245) from Thermo Fisher.

## Acknowledgements

M. Martínez for lentivirus plasmids and J. Roig for CRISPR plasmids. M. Robinson for sending the construct of the GST-γ-adaptin ear and M. Mapelli for the GST-LIC1 and LIC2 constructs. T. Surrey and C. Mitchel for sharing the porcine purified dynactin complex. This work was financed by BFU2017-82959-P to MIG, BES-2012-053341 to AB and BES-2015-071691 to IHP, from the Spanish government. The Andor Dragonfly 505 and the Zeiss LSM780 microscopes were funded by EQC2018-004541 EU FeDer and CSIC1501/18, respectively.

## Supplementary data

**Figure S1.**
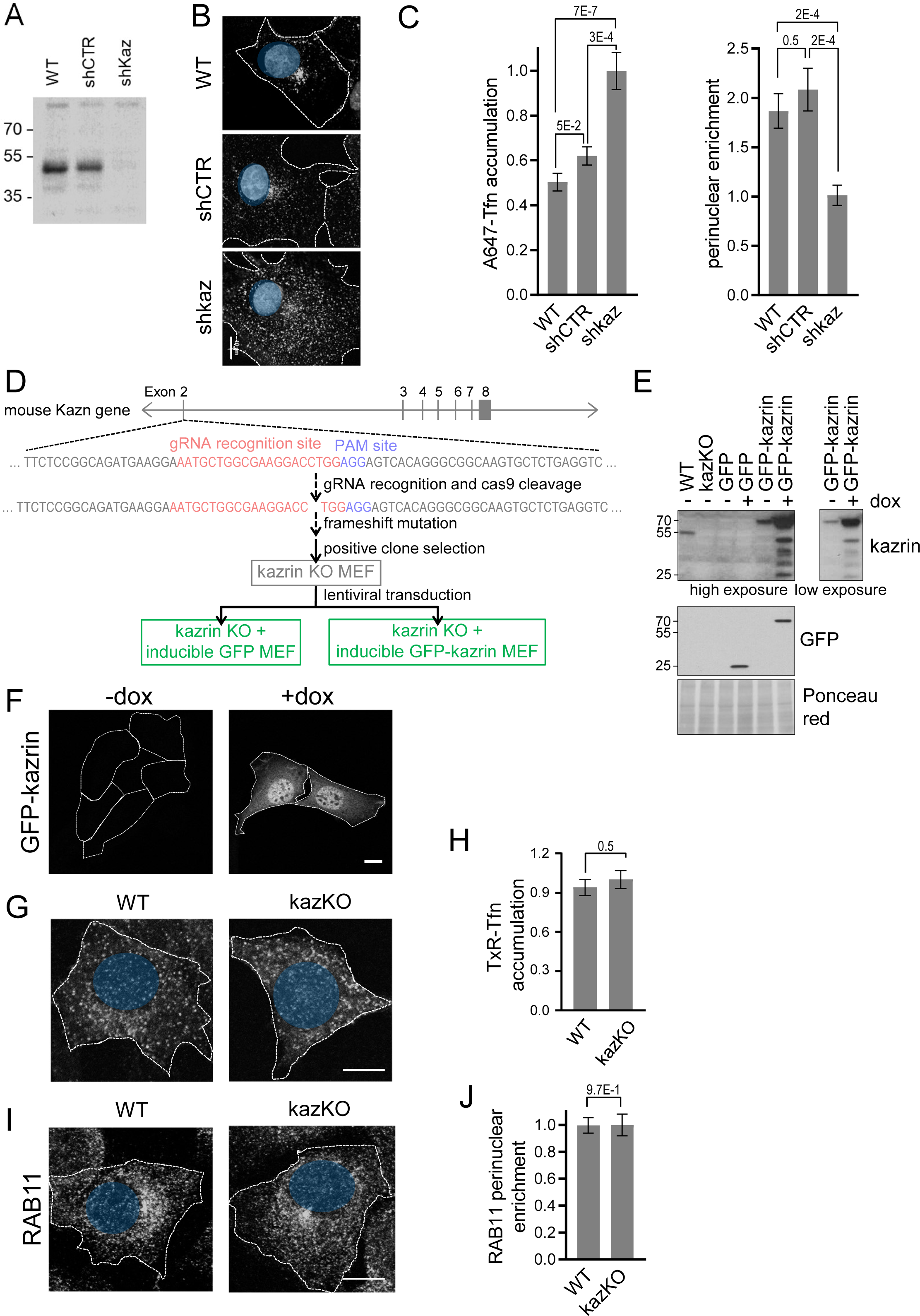
(**A**) Immunoblot against kazrin of protein extracts from Cos-7 cells either WT or expressing a non-target control shRNA (shCTR) or a shRNA against kazrin (shKaz). Proteins were extracted four days upon transduction and selection with puromycin. 18 μg of total protein extract was loaded per lane. The immunoblot was decorated with a polyclonal rabbit serum against the N-terminus of kazrin C. (**B**) Maximum intensity projections of confocal fluorescence micrographs showing either WT Cos-7 cells, or Cos-7 cell lines expressing a non-target control shRNA (shCTR) or a shRNA against kazrin (shKaz). Cells were exposed to Tfn-Alexa647 (A647-Tfn) for 2 hours before fixation. Dashed lines indicate the cell periphery. Bar, 10 µm. (**C**) Mean ± SEM of A647-Tfn fluorescence intensity accumulated per cell (left) or A647-Tfn perinuclear enrichment (right) after 2 hours incubation. For the A647-Tfn accumulation (left), the data was normalized to the mean value of the shkaz treated cells. p values of two-tailed Student t tests are shown. n > 25. For the perinuclear enrichment (right), the mean fluorescence intensity within a circle of 10 µm in the perinuclear region was quantified and divided by the total signal in the cell. The perinuclear enrichment data is normalized to the average of the shkaz treated cells. p values of two-tailed Student t tests are shown. n > 13 for each sample. (**D**) Strategy for the establishment of kazKO MEFs created with the CRISPR-Cas9 technology. The gRNA was designed to recognize a sequence at the beginning of exon 2 of the mouse Kazn gene, after the initiation codon of kazrin C, and followed by a PAM site. The CAS9 nuclease gene was transfected in a plasmid into the cells, together with the gRNA. CAS9 cleavage often leads to a frameshift mutation that impedes the expression of the gene. The plasmid encoding the gRNA and the CAS9 also encodes GFP, which allows sorting and isolation of transfected cells by FACS. The resulting clones were analysed by immunoblot, and those with no kazrin expression were selected. One of them was used as the base for another three cell lines in which genes encoding GFP or GFP-kazrin C were inserted in the genome. The inserted constructs were preceded by a tetracycline-response element. This was achieved by lentiviral transduction and selection by FACS. Thus, none of these cell lines have endogenous kazrin expression but expresses GFP or GFP-kazrin C upon doxycycline addition. (**E**) Immunoblots of cell lysates from WT and kazKO MEF or kazKO MEFs expressing GFP and GFP-kazrin C, in the presence (+) or absence (-) of 5 µg/ml doxycycline for 24 h (dox). The membranes were probed with a polyclonal rabbit serum against the N-terminus of kazrin C, α-GFP or stained with Ponceau red (as a loading control). A high and a low exposure for the kazrin signal are shown. (**F**) Confocal images of WT and kazKO MEF incubated for 10 minutes with 20 µg/ml TxR-Tfn at 37°C. Bar, 10 µm. (**G**) Mean ± SEM TxR-Tfn fluorescence intensity per cell for WT and kazKO cells. The fluorescence intensity in WT and kazKO cells was normalized to the mean value of the kazKO cells. The p value of a Mann Whitney test is shown. n > 100 for each sample. (**H**) Confocal images of GFP-kazrin C kazKO MEFs in the presence (+) or absence (-) of 5 µg/ml doxycycline for 24 h (dox). Scale bar, 10 µm. (**I**) Confocal images of WT and kazKO MEFs, fixed and stained with α-RAB11 and A488-conjugated secondary antibodies. The dashed lines indicate the cell periphery. The position of the nucleus is indicated in blue. (**J**) Mean ± SEM RAB11 perinuclear enrichment. The mean fluorescence intensity signal within a circle of 9 µm in the perinuclear region was quantified and divided by the RAB11 signal in the cell. The data was normalized to the mean of the kazKO cells. p values of two-tailed Student t tests are shown. n > 29 for each sample.

**Figure S2.**
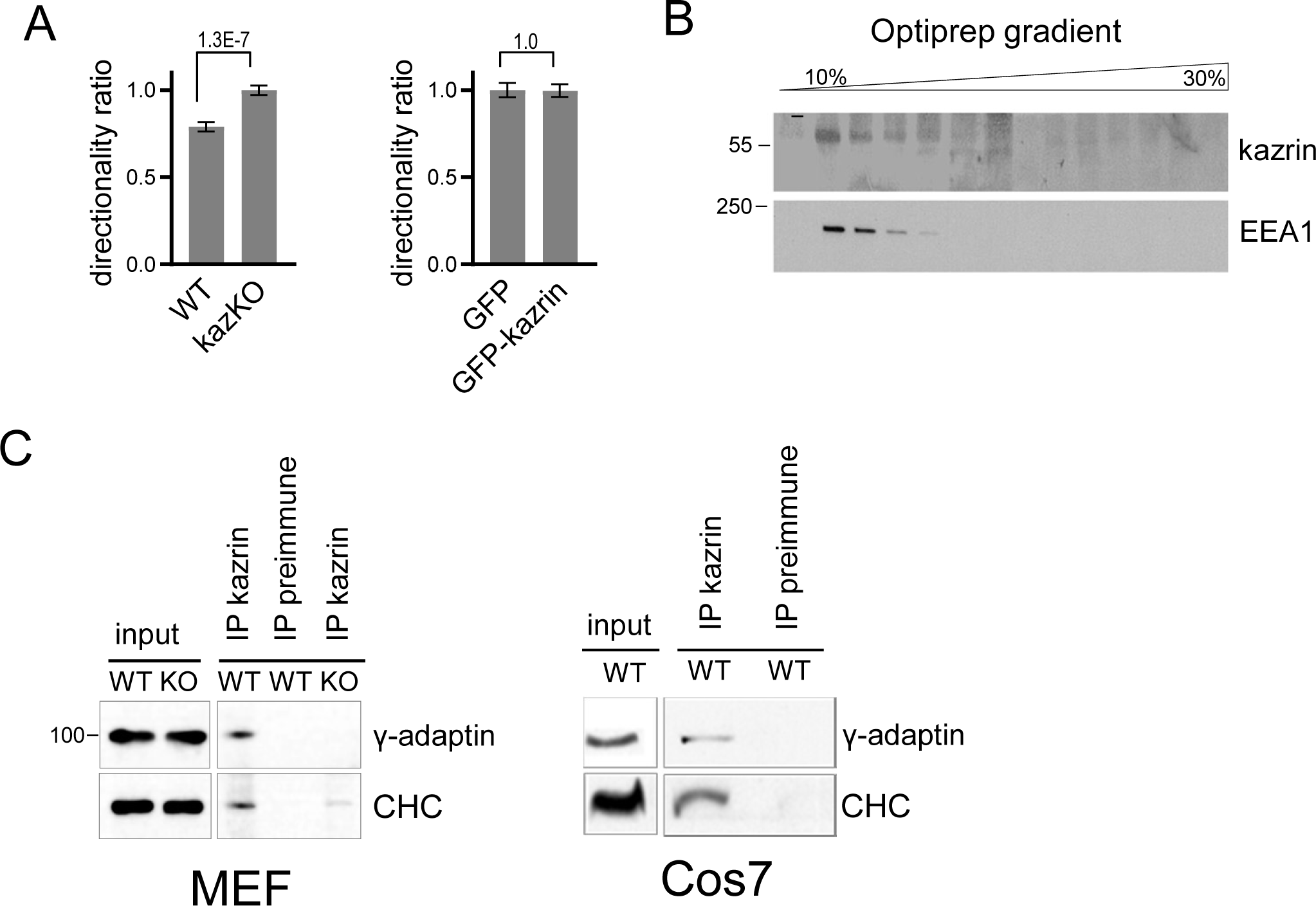
**(A)** Mean ± SEM directionality ratio of individually migrating WT and kazKO or kazKO MEF expressing low levels of GFP or GFP-kazrin C. The cells were embedded in Matrigel and tracked for 9 hours. The data was normalized to the corresponding kazKO cells. p values of two-tailed Mann-Whitney tests are shown. n > 155 per condition. (**B**) Immunoblots of an Optiprep density gradient fractionation of membrane lysates of Cos7 cells, probed with α-kazrin and α-EEA1 antibodies. (**C**) Immunoblot of protein A-Sepharose precipitates from WT or kazKO MEFs or Cos7 cells using a mixed serum against the N and the C-terminal portions of kazrin C or a pre-immunisation serum, probed with an α-γ-adaptin or α-Clathrin Heavy Chain (CHC) antibodies. Endogenous kazrin could not be detected with any of the tested antibodies in the immunoprecipitates because the antibody chain has a molecular weight similar to that of kazrin (approx. 50 Kda). kazKO MEF were used as specificity control instead.

**Figure S3.**
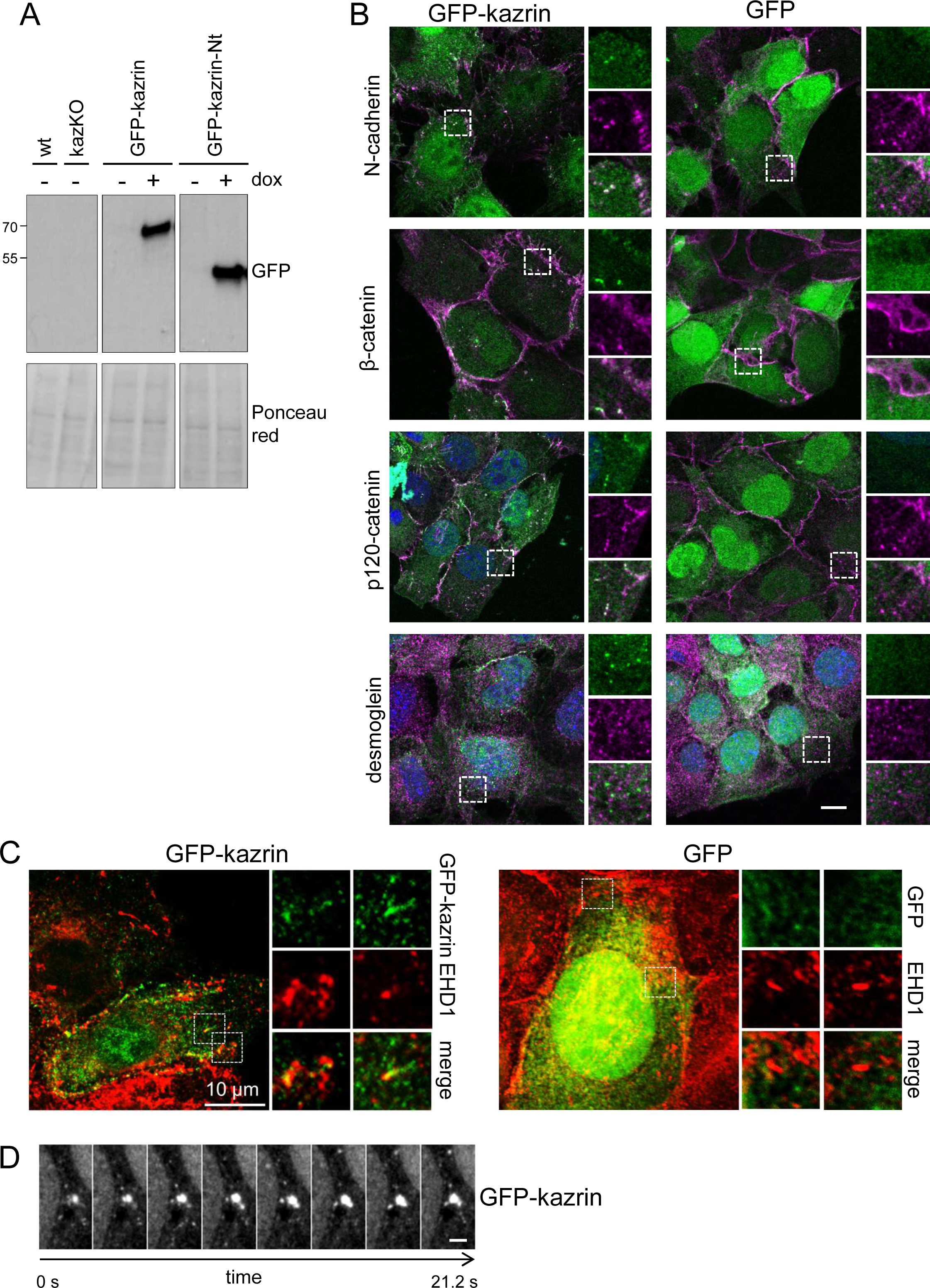
(**A**) Ponceau red staining (lower panels) and immunoblots against GFP of WT and kazKO MEFs, or kazKO cells expressing GFP-Kazrin C or a GFP-Kazrin C construct lacking the IDR (GFP-kazrin-Nt) in the absence (-) or presence (+) of of 5 µg/ml doxycycline for 24 h (dox). (**B**) Merged confocal images of kazKO MEFs expressing moderate levels of GFP or GFP-kazrin C, fixed and stained with α-N-cadherin, α-β-catenin, α-p120-catenin or α-desmogelin antibodies and adequate A568-conjugated secondary antibodies. Merged images and individual channels of 8x magnifications are shown. Scale bar, 10 μm. (**C**) Maximum intensity projections of merged confocal images of kazKO MEFs expressing moderate levels of GFP or GFP-kazrin C (GFP-kaz), fixed and stained with α-EHD1 or α-γ-adaptin and A568-conjugated secondary antibodies. Individual channels and merged images of 5 x 5 µm^2^ insets are shown on the right. (**D**) 2.65 second time-lapse confocal images of the perinuclear region of kazKO MEFs expressing moderate levels of GFP-kazrin C, Scale bar, 2 μm.

**Movie S1.** Videos of individually migrating WT and kazKO MEF, and kazKO MEF expressing low levels of GFP and GFP-kazrin C. The cells were embedded in Matrigel and imaged with an epifluorescence microscope.

**Movie S2.** Videos of dividing WT and kazKO MEF, and kazKO MEF expressing low levels of GFP and GFP-kazrin C, from the moment the mother cell attached to the substrate until the daughter cells were completely separated. The cells were embedded in Matrigel and imaged with an epifluorescence microscope.

**Movies S3 to S6.** 3D reconstructions Z stacks of kazKO MEF expressing moderate amounts of GFP-kazrin C loaded with TxR-Tfn at 16℃ to accumulate endocytic cargo in EEs. Cells were shifted to 37℃ and fixed after 10 minutes. The window is 5 x 5 µm^2^.

**Movies S7 to S10.** 3D reconstructions of Z stacks of kazKO MEF expressing moderate amounts of GFP-kazrin C, fixed and stained with α-EHD1 and A568-conjugated secondary antibodies. The window is 5 x 5 µm^2^.

**Movies S11 to S14.** 3D reconstructions of Z stacks of kazKO cells expressing moderate amounts of a GFP-kazrin C lacking the C-terminal predicted IDR (GFP-kazrin C-Nt) loaded with TxR-Tfn at 16℃ to accumulate endocytic cargo in EE. Cells were shifted to 37℃ and fixed after 10 minutes. The window is 5 x 5 µm^2^.

**Movies S15 & S16.** 3D reconstructions of Z stacks of kazKO cells expressing moderate amounts of GFP, loaded with TxR-Tfn at 16℃ to accumulate endocytic cargo in EE. Cells were shifted to 37℃ and fixed after 10 minutes. The window is 5 x 5 µm^2^.

**Movie S17.** 2.65 seconds time-lapse video of the perinuclear region of a kazKO MEF moderately expressing GFP-kazrin C (GFP-kaz) with a confocal microscopy. Scale bar, 2 μm.

**Movie S18.** 3 seconds time-lapse live-cell movies showing TxR-Tfn loaded endosomal dynamics in WT MEF. The window is 10 x 10 µm^2^. Cells were loaded with TxR-Tfn at 16℃ to accumulate endocytic cargo in EEs and imaged immediately after shift to 37℃. The image corresponds to the maximum intensity Z projection.

**Movie S19.** 3 seconds time-lapse live-cell movies showing TxR-Tfn loaded endosomal dynamics in kazKO MEF. The window is 10 x 10 µm^2^. Cells were loaded with TxR-Tfn at 16℃ to accumulate endocytic cargo in EEs and imaged immediately after shift to 37℃. The image corresponds to the maximum intensity Z projection.

**Movie S20.** 3 seconds time-lapse live-cell movies showing TxR-Tfn loaded endosomal dynamics in kazKO MEF expressing low levels of GFP. The window is 10 x 10 µm^2^. Cells were loaded with TxR-Tfn at 16℃ to accumulate endocytic cargo in EEs and imaged immediately after shift to 37℃. The image corresponds to the maximum intensity Z projection.

**Movie S21.** 3 seconds time-lapse live-cell movies showing TxR-Tfn loaded endosomal dynamics in kazKO MEF expressing low levels of GFP-kazrin C. The window is 10 x 10 µm^2^. Cells were loaded with TxR-Tfn at 16℃ to accumulate endocytic cargo in EEs and imaged immediately after shift to 37℃. The image corresponds to the maximum intensity Z projection.

**Movie S22.** 3 seconds time-lapse live-cell movies showing TxR-Tfn loaded endosomal dynamics in kazKO MEF expressing low levels of a GFP-kazrin C construct lacking the C-terminal predicted IDR (GFP-kazrin c-Nt). The window is 10 x 10 µm^2^. Cells were loaded with TxR-Tfn at 16℃ to accumulate endocytic cargo in EEs and imaged immediately after shift to 37℃. The image corresponds to the maximum intensity Z projection.

